# APP substrate ectodomain defines Aβ length by restraining γ-secretase processivity and facilitating product release

**DOI:** 10.1101/2023.09.13.557360

**Authors:** Matthias Koch, Thomas Enzlein, Shu-Yu Chen, Dieter Petit, Sam Lismont, Martin Zacharias, Carsten Hopf, Lucía Chávez-Gutiérrez

## Abstract

Sequential proteolysis of the amyloid precursor protein (APP) by γ-secretases (GSECs) generates amyloid-β (Aβ) and defines the proportion of short-to-long Aβ peptides, which is tightly connected to Alzheimer’s disease (AD) pathogenesis.

Here, we study the mechanism controlling substrate processing by GSECs and defining product length. We found that polar interactions established by the APP_C99_ ectodomain (ECD), involving but not limited to its juxtamembrane region, restrain both the extent and degree of GSEC processive cleavage by destabilizing enzyme-substrate (E-S) interactions. We show that increasing hydrophobicity at APP_C99_-ECD – due to mutation or ligand binding – attenuates this substrate-driven product release mechanism, and rescues the effects that AD pathogenic variants exert on Aβ profiles. In addition, our study reveals that APP_C99_-ECD facilitates the paradoxical production of longer Aβs caused by some GSEC inhibitors that act as high-affinity competitors to the substrate.

These findings assign a pivotal role to the substrate ECD in the sequential proteolysis by GSEC and suggest it as a sweet spot for the potential design of APP targeting compounds selectively promoting its processing by GSEC.

## Introduction

γ-Secretases (GSECs) are multifaceted proteolytic switches controlling numerous signalling processes in pathophysiology (Chávez-Gutiérrez & Szaruga, 2020; Jurisch-Yaksi *et al*, 2013). They are multimeric, membrane-embedded complexes that consist of nicastrin (NCSTN), anterior pharynx defective 1 (APH1), presenilin enhancer 2 (PEN-2) and presenilin (PSEN, catalytic subunit) (Hur, 2022). GSECs recognize type I membrane proteins presenting short (<50 aa) ectodomains (Güner *et al*, 2020; Struhl & Adachi, 2000; Funamoto *et al*, 2013) and cleave them sequentially within their transmembrane domains (TMD) in a process referred to as regulated intramembrane proteolysis (RIP) (Brown *et al*, 2000). An initial (endopeptidase) ε-cleavage may switch on/off signalling cascades at the membrane by either releasing soluble courier proteins intracellularly or destroying membrane-embedded signalling proteins. The release of the Notch intracellular domain (NICD) from the membrane (Schroeter *et al*, 1998; De Strooper *et al*, 1999), a key event in the Notch-signalling pathway, illustrates the former; while cleavage of the ‘pro-apoptotic signalling’ C-terminal fragments of the p75 neurotrophin receptor (p75-CTF) exemplifies the latter case (Franco *et al*, 2021).

Two structures of the initial enzyme-substrate (E-S) complexes with either the amyloid precursor protein (APP) (**Fig 1A**) or Notch are available. Despite the low sequence similarity of these substrates, the structures show a remarkable similar overall E-S conformation (Zhou *et al*, 2019; Yang *et al*, 2019), wherein PSEN embraces the (mainly) helical TMD of the substrate. Near the active site, PSEN and the substrate form a hybrid (E-S) β-sheet structure which exposes the scissile bond to the catalytic Asp dyad; thus, facilitating the ε-cleavage and consequent release of the intracellular domain of the substrate. The remaining TMD stub is then sequentially cut (carboxypeptidase-like γ-cleavages), ultimately resulting in the release of an N-terminal peptide (**Fig 1B**).

**Figure 1.**
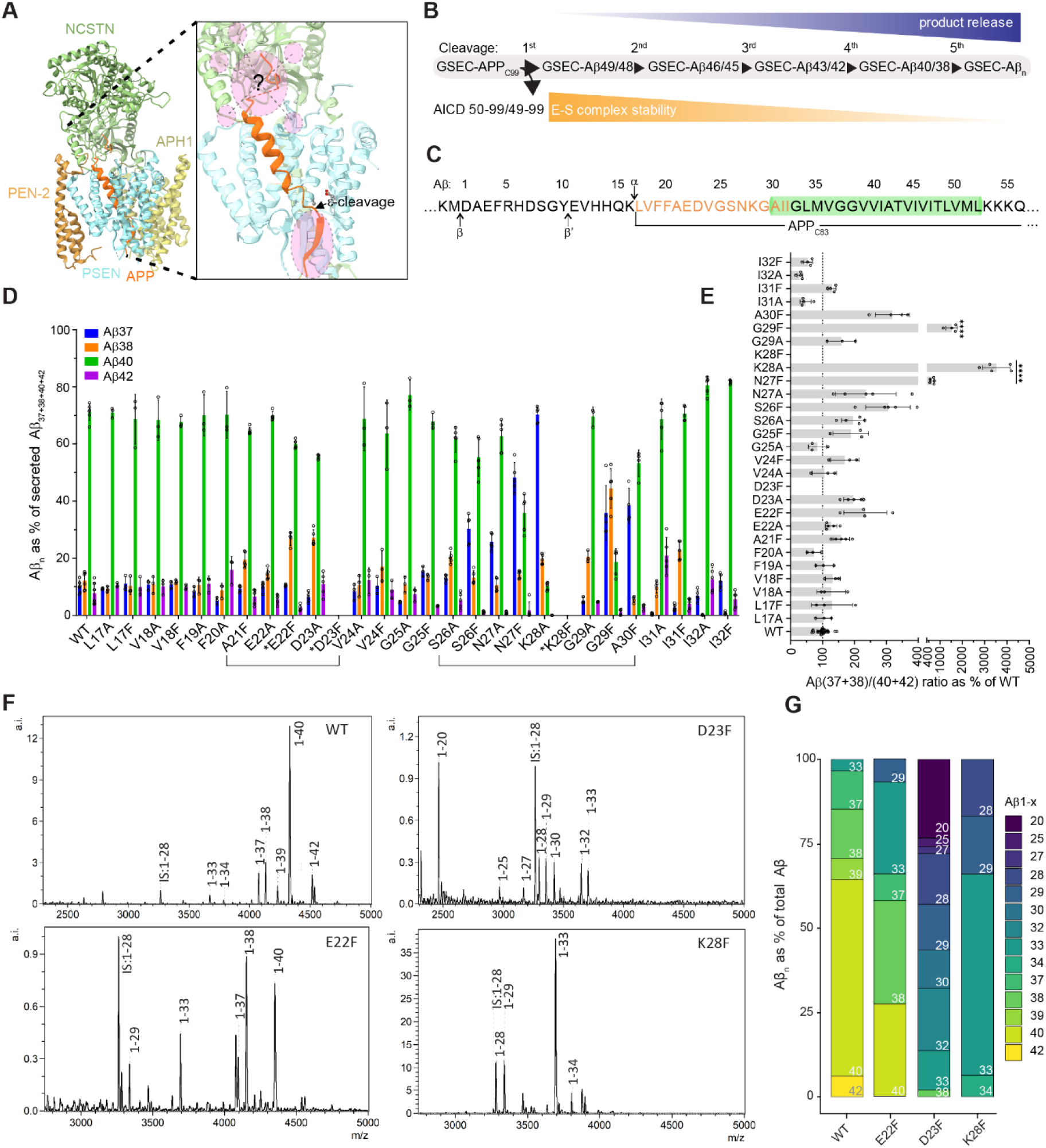
Delineation of sequence determinants in APP_C99_-ECD modulating GSEC processivity. (A) GSEC-APP_C83_ complex (PDB: 6IYC) shows the substrate TMD C-terminally unwound and interacting with PSEN1 via an induced hybrid β-sheet. N- and C-terminal E-S interactions are highlighted in purple; N-terminal interactions play a crucial, yet still elusive role in GSEC-proteolysis. (B) Schematic representation of the sequential cleavage of APP_C99_ by GSEC. Each cleavage decreases E-S stability and increases product release probability. (C) APP_C99_ (1-56 aa) sequence with APP-TMD highlighted in green and residues that were subjected to Ala/Phe mutagenesis in orange. BACE (β, β’) and ADAM10 (α) cleavage sites are indicated. (D) Aβ peptides secreted by HEK293T cells transiently overexpressing WT or mutant (Ala/Phe) APP_C99_ substrates were quantified by multiplex MSD ELISA. Aβ peptides are shown as % of the sum of all measured peptides (profiles). The A21-D23 and S26-A30 stretches are underlined to highlight their critical roles in GSEC processivity. Mean ± SD; N ≥ 3 independent experiments. Generation of Aβ peptides from mutant substrates marked with an asterisk (*) was drastically decreased and Aβ profiles (except E22F) were not determined due to low Aβ signals in ELISA. (E) Aβ(37+38)/(40+42) ratios informed about mutation-driven changes in GSEC processivity. One-way ANOVA followed by Dunnett’s post hoc test with comparison to WT was used to determine statistical significance (P < 0.05). ****P < 0.0001 F (DFn, DFd): F (26, 103) = 127.1. (F) Representative MALDI-TOF MS spectra of Aβs IPed from CM of HEK293T cells expressing WT or mutant APP_C99_ (E22F, D23F, K28F). Synthetic Aβ1-28 peptide was used as internal standard (IS). (G) Aβ profiles determined from data presented in (F). Aβ peptides that appeared in at least two independent biological replicates are shown. Mean; N ≥ 3 independent experiments.

The generation of N-terminal amyloid-β (Aβ) peptides from APP by GSECs plays a pivotal pathogenic role in Alzheimer’s disease (AD) (Selkoe & Hardy, 2016), and therefore has been a matter of intense research. Removal of the APP ectodomain (ECD) by β-secretase (BACE) generates a membrane-embedded APP C-terminal fragment of 99 amino acids (aa) in length (APP_C99_) (Vassar *et al*, 1999), which is then cleaved in a stepwise manner (processivity) by GSECs along two major product lines (Takami *et al*, 2009; Matsumura *et al*, 2014) (**Fig 1B**).

Mutations in *PSEN1/2* and *APP* causing familial AD (FAD) destabilize E-S interactions (Szaruga *et al*, 2017) and thereby promote the release of partially digested, longer Aβ peptides such as Aβ42 (Szaruga *et al*, 2015), Aβ43 (Kretner *et al*, 2016; Saito *et al*, 2011; Veugelen *et al*, 2016), and possibly Aβ45 and Aβ46 (Liu *et al*, 2017). Pathogenic mutations consistently shift the Aβ product profile towards longer Aβ peptides (Szaruga *et al*, 2015), and the short-to-long Aβ(37+38+40)/(42+43) ratio linearly correlates with the age at clinical AD onset (AAO) in carriers of PSEN1 variants (Petit *et al*, 2022a). The pathological relevance of shifts in the proportion of short-to-long Aβ peptides has also been demonstrated in the sporadic and most common form of AD (SAD) (Liu *et al*, 2022), where changes in the Aβ37/42 ratio distinguish AD patients from cognitively normal subjects. In both familial and sporadic AD, increments in longer Aβs (≥ 42 aa) are proposed to promote the assembly of (yet-to-be-identified) toxic Aβ assemblies that initiate molecular and cellular cascades leading to neurodegeneration (Hardy & Higgins, 1992; Selkoe & Hardy, 2016).

Allosteric GSEC inhibitors (GSIs), such as DAPT (Qi-Takahara *et al*, 2005; Yagishita *et al*, 2006) and semagacestat (Tagami *et al*, 2017), also lead to increases in the production of longer Aβ peptides, while partially inhibiting GSEC activity. This paradoxical ‘FAD-mimicking’ effect on Aβ production has been postulated to contribute to the worsening of cognition observed in AD patients treated with the GSI semagacestat (Tagami *et al*, 2017). The underlying mechanism(s) remain unclear.

Genetic and clinical observations thus assign high pathophysiological relevance to the sequential cleavage of APP by GSEC. Nevertheless, its molecular underpinnings are poorly understood. Here, we investigate the mechanisms controlling the processive proteolysis of APP by GSECs that define Aβ product length. Our investigations show that the ECD of the substrate (APP_C99_/Aβ) restricts processivity as well as markedly destabilizes E-S interactions and pinpoint polar interactions involving but not limited to its juxtamembrane region as the critical underlying feature. We found that increased hydrophobicity in APP_C99_-ECD, due to substitutions at key Lys28 and Asp23 positions, mitigates APP_C99_-ECD-driven destabilization and markedly promotes efficient and extended sequential proteolysis of Aβ. Notably, attenuation of this substrate-driven product-release mechanism rescues the effects of AD pathogenic variants in GSEC and APP on Aβ profiles.

Our analyses of GSEC-mediated Notch cleavage indicate that this substrate-driven mechanism is of general application. Collectively, these data support a model in which polar/charged residues in the membrane-proximate region in the substrate ECD form an energy barrier that serves as a ‘pivot’ in the product release mechanism, and suggest that stochastic overcoming of this barrier facilitates further threading of the substrate. Finally, data on the inhibition of GSEC by DAPT and semagacestat show that these inhibitors act as high-affinity competitors to substrates, and that the destabilizing nature of APP_C99_-ECD facilitates the GSI’s paradoxical effects on the production of longer Aβ peptides.

Our studies assign a pivotal role to the substrate’s ECD in the sequential proteolysis of GSEC and suggest a potential sweet spot for the design of compounds selectively promoting efficient APP processing.

## Results

### Sequence determinants in APP_C99_-ECD, involving but not restricted to residues Asp23 and Lys28, control GSEC processivity

The efficiency of the sequential proteolysis of APP by GSECs plays a pivotal role in AD pathogenesis and its modulation by small compounds is being pursued as a therapeutic strategy (Luo & Li, 2022). Previous studies have shown that mutations in the APP substrate modulate its processing by GSEC to promote the generation of shorter Aβ peptides. Most importantly, the literature highlights a remarkable link between Lys28 situated in the juxtamembrane region of APP_C99_ (**Fig 1A, upper purple circle**) and Aβ product length, with substitutions to Ala or Glu markedly shifting production towards the generation of very short Aβ34 and Aβ33 peptides (Petit *et al*, 2019; Kukar *et al*, 2011; Jung *et al*, 2014; Page *et al*, 2010; Ousson *et al*, 2013). These findings support the notion that intrinsic – yet to be defined – determinants in the substrate play a major role in defining enzyme processivity and thereby Aβ product length. To gain insights into the underlying mechanism(s), we fist mapped the sequence determinants in APP_C99_-ECD that modulate the sequential γ-cleavages by performing an alanine/phenylalanine (Ala/Phe) mutagenic scan of the Leu17-Ile32 stretch (**Fig 1C)**. This region corresponds to the ECD of the shorter APP_C83_ substrate (generated by ADAM10), which contains all features required for efficient GSEC-mediated proteolysis (Funamoto *et al*, 2013). We transiently expressed WT or mutant APP_C99_ in HEK293T cells, and measured the levels of secreted Aβ37/38/40/42 peptides in the conditioned media (CM) by ELISA. We found that the K28A, K28F, D23F and E22F mutations drastically lowered all measured peptides (**Fig EV1A**), despite robust expression levels of WT/mutant substrates (**Fig EV1B**). These peptides, from now on referred to as canonical Aβs, were also significantly decreased by the A21F-, D23A-, G25F, N27F, G29F-, A30F-, I31F- and I32F mutations (**Fig EV1A**), and increased by the F20A substitution in APP_C99_. The latter effect is consistent with previous findings showing that Phe20 is part of an inhibitory domain (Aβ17-23) in APP/Aβ and that disruption of this domain elevates GSEC activity (Tian *et al*, 2010).

Aβ profiles, estimated as the percentage of each peptide relative to the total canonical Aβ levels, were analysed for all substrates, except the K28F and D23F mutants due to very low peptide levels. Aβ profiles revealed marked shifts towards enhanced generation of short Aβ37 and Aβ38 peptides for Phe substitutions at positions Ser26-Ala30, while the analogous mutations to Ala displayed similar but milder effects (**Figs 1D and EV1C**). In addition, Ala substitutions at positions Ala21-Asp23 caused mild increments in Aβ38 (**Fig 1D**).

To estimate the efficiency of the sequential processing, we determined the Aβ(37+38)/(40+42) ratio (Chávez-Gutiérrez *et al*, 2012). The data revealed 6-, 30- and 15-fold increases in GSEC processivity for the mutant N27F-, K28A- and G29F- APP_C99_ substrates, respectively (**Fig 1E**).

We next investigated the processing of mutant K28F and D23F substrates in a cell-free GSEC activity assay, using purified enzyme and substrate. We found no significant differences in AICD product levels generated from WT, K28F or D23F mutants, relative to the (reference) D23A, K28A, S26F, N27A/F mutant APP_C99_ substrates (**Fig EV1D**). These results imply that the lack of generation of canonical Aβ peptides resulted from mutation-driven changes in processivity, rather than from changes in overall substrate endopeptidase processing. To further investigate this, we immunoprecipitated (IPed) secreted Aβ peptides from CM and analysed them by MALDI-TOF mass spectrometry. We included the E22F substrate in this analysis given its profound effects on canonical Aβ levels (**Fig EV1A**). As expected, mass spectra showed Aβ40 as the major product generated from the from the WT substrate, and lower amounts of the Aβ33, Aβ37, Aβ38, Aβ39 and Aβ42 peptides. Treatment with the GSEC inhibitor L-685,458 (InhX) completely abolished Aβ production (**Fig EV1E**) demonstrating that the detected peptides were produced in a GSEC-dependent manner. This analysis also revealed Aβs varying from 29 aa to 40 aa and from 20 aa to 38 aa in lengths for the E22F and D23F mutant substrates, respectively. Moreover, Aβ33 and minor amounts of Aβ34, Aβ29 and Aβ28 were measured for the K28F mutant substrate (**Figs 1F-G**). These data support the implication of the substrate APP-ECD in the modulation of the sequential GSEC-mediated proteolysis, and pointed at increased hydrophobicity in the Ala21-Asp23 and S26-Ala30 regions as the critical feature in the modulation of Aβ product length, with the Lys28 and Asp23-Glu22 positions as key determinants of the sequential cleavage mechanism.

### Direct and linear relationship between hydrophobicity at position 28 in APP_C99_**-**ECD and GSEC processivity

The prominent role of position APP_C99_-28 in the modulation of Aβ processing let us to evaluate the effects of different aminoacidic substitutions at this site on the secreted Aβ pool. We quantified both canonical and total Aβ levels in CM of HEK293T cells overexpressing WT or mutant APP_C99_ substrates. Total Aβ levels measured by ELISA with the 4G8 and 6E10 anti-Aβ antibodies, as capturing and detection reagents, were used as proxy in the estimation of global endopeptidase activity.

Total Aβ, normalized to substrate expression (**Fig EV2A**), demonstrated significant increases for 13 out of 19 substrates, and similar levels for the K28R, K28Y, K28W, K28F, K28L, K28I mutants, relative to WT (**Fig 2A**). In contrast, all substitutions drastically reduced the summation of secreted canonical Aβs, with the exception of K28R (**Figs 2B and EV2B**). The mismatch between total and canonical Aβ levels implied substantial shifts in Aβ product profiles for most of the tested mutant substrates. In addition, our analysis revealed a connection between the levels of canonical Aβs and the hydrophobicity of the substituted side chain. Note that **Figures 2A-B** present the aa substitutions according to the Kyte-Doolittle (K-D) hydrophobicity scale (**Fig 2C**) (Kyte & Doolittle, 1982).

**Figure 2.**
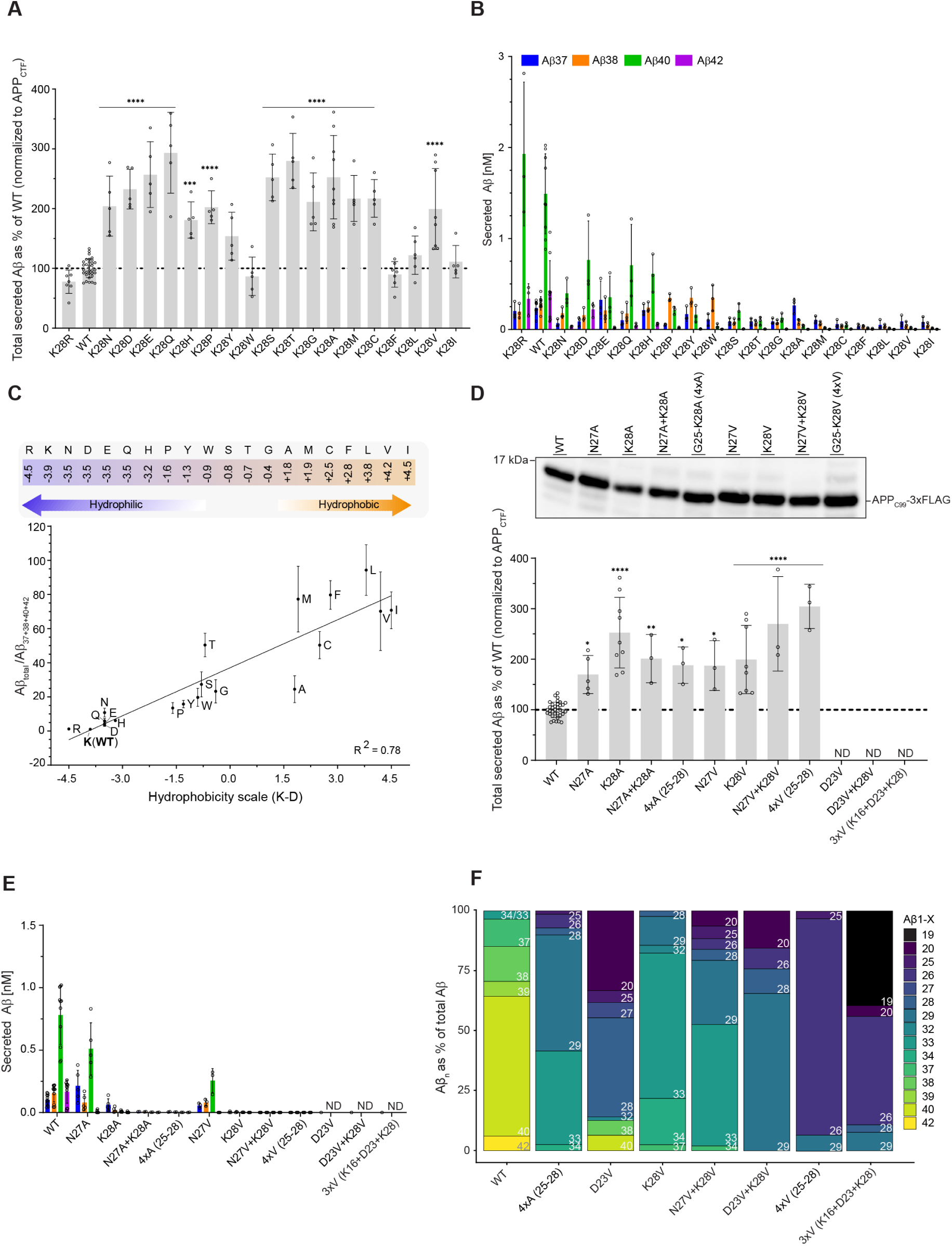
Hydrophobicity at APP_C99_-28 increases sequential γ-cuts on Aβ. (A) Total secreted Aβ levels generated by HEK293T cells expressing WT- or K28X-APP_C99_ mutants were analysed by ELISA. In this assay, we used anti-APP 4G8 and 6E10 antibodies as capturing and detection reagents, respectively. Data was normalized to APP_C99_ expression levels determined by western blot (**Figure S2A**). Mean ± SD; N ≥ 5 independent experiments. One-way ANOVA followed by Dunnett’s post hoc test was used to determine statistical significance (P < 0.05) relative to WT. ****P < 0.0001, ***P < 0.001 F (DFn, DFd): F (19, 119) = 22.25. (B) Secreted Aβ37/38/40/42 peptides analysed by multiplex MSD ELISA. Mean ± SD; N = 3 independent experiments. (C) The K-D scale (Kyte & Doolittle, 1982) is shown in the upper panel. The Aβ_total_/Aβ(37+38+40+42) ratio, used as an estimate of GSEC processivity, shows a positive correlation with hydrophobicity at position APP_C99_-28 (lower panel). The WT ratio was set to 1; values > 1 thus indicate increased GSEC processivity. Mean ± SD; N = 3 independent experiments. R^2^=0.78; Y = 9.377*X + 37.07. (D) Total secreted Aβ of HEK293T cells expressing either WT APP_C99_ or single, double, triple and quadruple alanine or valine substitutions at indicated positions were analysed by ELISA. Mean ± SD; N ≥ 3 independent experiments. One-way ANOVA followed by Dunnett’s post hoc test with comparison to WT was used to determine statistical significance (P < 0.05). ****P < 0.0001, ***P < 0.001, **P < 0.01, *P < 0.1 F (DFn, DFd): F (11, 63) = 19.51. (E) Aβ(37,38,40,42) peptide levels were analysed in CM from cells expressing WT/mutant APP substrates by multiplex ELISA. (F) Aβ profiles determined by (MALDI-TOF) mass spectrometry-based analysis of WT and mutant Aβ peptides IPed from CM collected from cells expressing selected APP_C99_ mutants. Data shown as mean N ≥ 3 independent experiments with exception of K28V, N = 2 independent experiments. *ND = not determined; substitutions at position APP_C99_-23 disrupt the epitope of the anti-Aβ (4G8) antibody used to quantify the total peptide pool by ELISA, thus analysis was only performed by MALDI-TOF MS.

As the total Aβ pool (Aβ_total_) contains shorter peptides in addition to the canonical Aβs, we reasoned that the total-to-canonical Aβ peptide ratio (Aβ_total_/Aβ(37+38+40+42)) provides an estimate of the degree of GSEC-mediated processivity on WT or mutant APP substrates. Assessment of this ratio showed that hydrophobic substitutions substantially increased the degree of processing of mutant substrates (**Fig S2C**), and revealed a significant linear correlation between GSEC processivity and hydrophobicity at position 28 in APP_C99_-ECD (**Fig 2C**). Our analysis thus shows that hydrophobicity at the juxtamembrane region of APP_C99_ is a key determinant governing both cleavage efficiency and extent of Aβ processing by GSEC.

### Increased hydrophobicity in APP_C99_-ECD promotes substrate threading and extends processing by GSEC

We next investigated potential additive and/or synergistic effects between APP_C99_-28 and neighbouring (Ser26, Asn27 or Asp23) positions in the modulation of Aβ processing by GSEC. Specifically, we asked whether hydrophobicity and/or helicity in APP_C99_-ECD influence enzyme processivity. We mutated 2 aa (D23+K28 or N27+K28) and 4 aa stretches (G25-K28) in APP to Val or Ala, and analysed Aβ product levels secreted from HEK293T cells overexpressing the respective substrates (**Fig 2D)**. Single mutations at Asp23, Asn27 and Lys28 were used as references. Of note, both Val and Ala have high hydrophobicity indexes (Kyte & Doolittle, 1982; Pace & Scholtz, 1998) but, in contrast to Val, Ala has a high helical propensity (Gregoret & Sauer, 1998).

We found significant increases in total Aβs and drastic reductions in canonical Aβ levels for the double and quadruple mutations (**Figs 2D and 2E, respectively**). Further, mass spectrometry-based analysis of IPed Aβs generated from the D23V, K28V, D23V+K28V, N27V+K28V, 4xV (25-28) and 4xA (25-28) mutated substrates (**Figs 2F and EV2D**) showed that very similar peptide profiles were generated from the 4xA and N27V+K28V APP_C99_ substrates, with Aβ33 and Aβ29 as the main products and Aβ34 and shorter (≤ 28 aa) peptides as minor ones. Aβ profiles generated from the double D23V+K28V vs single D23V or K28V mutant substrates revealed that even shorter peptides (29 aa - 20 aa) were generated from the double mutant, relative to the single ones. In addition, a substantial shift towards shorter Aβ peptides, with Aβ26 as the predominant product, was observed for the 4xV substrate. Of note, the generation of the very short Aβ29-Aβ20 peptides imply extended substrate threading, with a part of APP_C99_-ECD going into the membrane-embedded catalytic pore in PSEN. The next positively charged residue N-terminal to Lys28 in the D23V+K28V substrate is Lys16 (**Fig 1C**). To investigate whether this residue restricts further processing of the D23V+K28V substrate, we additionally substituted it to Val. While Aβ26 still represented the main product generated from this 3xV (16+23+28) mutant substrate (**Figs 2F and EV2D**), the even shorter Aβ19 accounted for about 40% of the total Aβ products. In conclusion, the mutation-driven shifts on Aβ processing (N27V+K28V < 4xA < D23V+K28V < 4xV < 3xV) show cooperative effects between the tested hydrophobic substitutions and indicate that hydrophobicity, rather than helicity (see also **Fig EV2E**), in APP_C99_-ECD is a critical feature in modulating Aβ peptide length.

### Notch-His17 – as Lys28 in APP_C99_ – restrains substrate threading and sequential GSEC processing

We then evaluated whether this substrate-driven mechanism also applies to the cleavage of Notch, a GSEC substrate well-known for its pivotal roles in cell differentiation and proliferation (Jurisch-Yaksi *et al*, 2013). We selected His17 in Notch (**Fig 3A**) for mutagenic analysis since structural data for the GSEC-Notch complex (PDB: 6IDF (Yang *et al*, 2019)) shows this residue at the juxtamembrane region of the substrate (**Figs 3A-B**); thus, suggesting that it could play a role similar to that of Lys28 in APP_C99_. We mutated His17 to either Gln, Phe, Lys or Asp (H17Q, H17F, H17K or H17D) in a Notch-based substrate (**Figs 3A**), expressed WT and mutant (HA-tagged) substrates in HEK293T cells and IPed secreted Notch N-terminal fragments (Nβ) using an anti-HA antibody. Importantly, mass spectrometry analysis of the (purified) Notch substrate indicated the presence of three substrate lengths due to imprecise signal peptide (SP) cleavage (**Figs 3A and 3C**). The observed distinct mass peaks were however assigned to derived Nβ products with high mass accuracy (**Figs EV3A**). Consistent with previous findings (Wanngren *et al*, 2012; Okochi *et al*, 2002, 2006), Nβ peptides with lengths ranging between 17 aa and 27 aa were generated from the WT Notch substrate (**Figs 3D, top panel and 3E**). The analysis of mutant product profiles showed that longer Nβ26/27/29 fragments were generated from the Notch-H17K substrate, relative to the WT. The introduction of a negatively charged aa at this position (H17D) had relatively small overall effects on Nβ processing; enhancing the generation of Nβ25/23, while lowering Nβ26, and supressing the generation of the minor Nβ22 and Nβ17 products. Furthermore, substitution of His17 by the polar, not charged Gln (H17Q), mildly increased processivity, while its replacement with a hydrophobic aa (H17F) strongly extended Nβ processing (**Figs 3D-E).** These data support a general model for GSEC processivity wherein the hydrophobic nature of the juxtamembrane region of the substrate (and ECD, as demonstrated above) critically modulates the efficiency and extent of the sequential cleavage mechanism.

**Figure 3.**
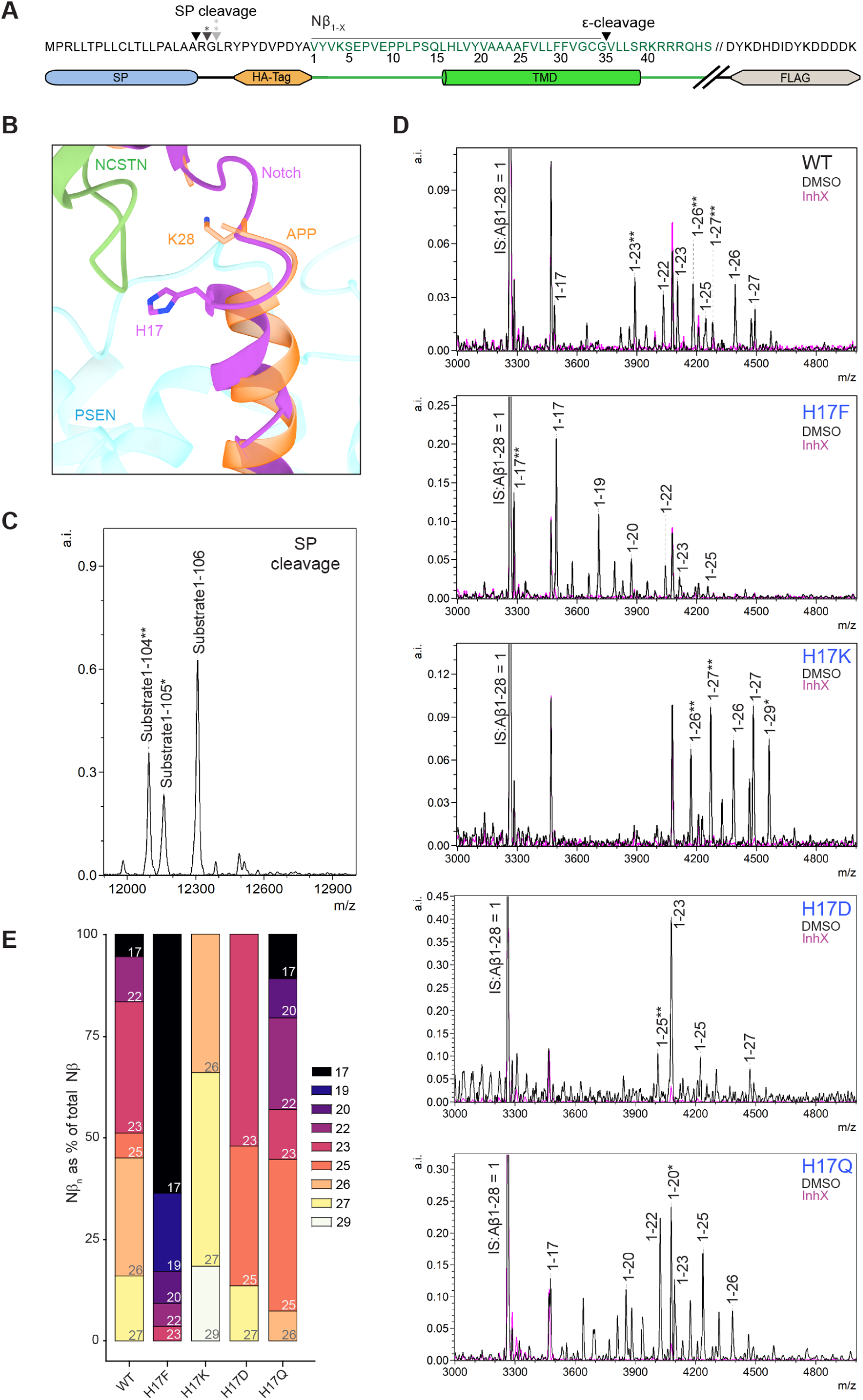
Notch sequential proteolysis is modulated by His17. (A) Overview of the mouse Notch construct used in this study. (B) Superimposition of high-resolution maps of GSEC bound to APP_C83_ (PDB: 6IYC) and Notch (PDB: 6IDF) shows positions Lys28 and His17 in APP and Notch, respectively. (C) Mass spectra of WT Notch purified through its 3x-FLAG-tag from HEK293T cells reveal generation of distinct substrates due to imprecise signal peptide (SP) cleavage. (D) HA-Nβs generated from WT and mutant (H17F, H17K, H17D, H17Q; depicted from top to bottom) Notch substrates were IPed from CM of HEK293T and analysed by MALDI-TOF MS. Representative mass spectra are shown; Notch-WT and Notch-H17F, N = 3; Notch-H17K, -H17D and -H17Q, N = 2 independent experiments. The different substrates (Figure 3C) and additional N-terminal HA-tag were considered for mass assignment (**Figure EV3A**). Most signal peaks were assigned and GSI treatment (data in purple) demonstrate that fragments are generated in a GSEC- dependent manner. (E) Nβ profiles determined by mass spectrometry-based analysis of WT/mutant peptides IPed from CM from cells expressing the indicated Notch substrates. Mean values of Nβs are shown, which appeared in at least two independent experiments.

### APP_C99_ Lys28 undermines GSEC-Aβ complex stability

Our previous studies have shown that factors destabilizing or stabilizing GSEC-Aβ_n_ interactions promote the generation of longer or shorter Aβ peptides, respectively (Szaruga *et al*, 2017; Petit *et al*, 2022c). We therefore investigated whether hydrophobicity at position 28 in APP_C99_ increases the production of shorter Aβs by stabilizing E-S interactions. We reasoned that mutations in APP, if increasing GSEC-Aβ_n_ stability, would counteract the detrimental effects that detergent and/or elevated temperature exert on these complexes (Szaruga *et al*, 2017; Petit *et al*, 2019). To assess this possibility, we incubated purified GSEC and WT or mutant (K28A, K28F) APP_C99_-3xFLAG substrates over a temperature gradient (37°C - 55°C) and determined Aβ product profiles by mass spectrometry (**Figs 4A-B**). These thermoactivity assays have been proven to be informative for the assessment of the effects of mutations and or ligands on the stability of GSEC-APP/Aβ_n_ interactions (Szaruga *et al*, 2017; Petit *et al*, 2019). Consistent with previous analyses (Szaruga *et al*, 2017), detergent-solubilization per se destabilized E-S interactions, thus enhancing the production of longer Aβ42-46 peptides from WT APP_C99_ at 37°C, relative to native conditions **(Figs 4A and EV4A vs 1G**); and further (thermal) destabilization shifted profiles towards production of even longer Aβ43-48 peptides (**Figs 4A (37°C vs 55°C) and 4B**).

**Figure 4.**
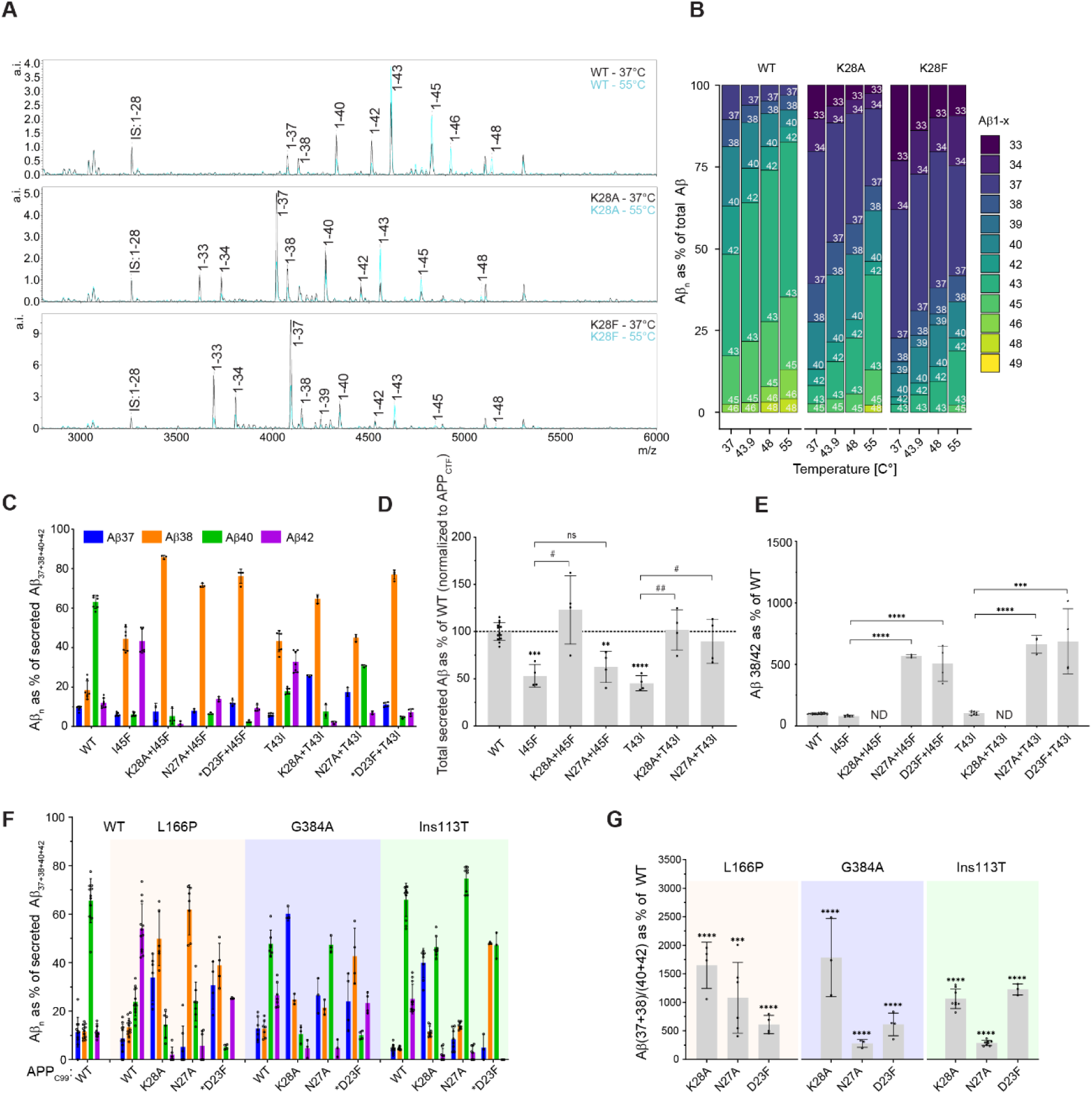
Increased hydrophobicity in APP_C99_-ECD rescues FAD-linked alterations in APP processing. (A) Cell-free activity assays using purified GSEC (PSEN1, APH1B) and WT or mutant (K28A, K28F) APP_C99_-3xFLAG performed at 37°C (black) or 55°C (blue). Aβ products were determined by MALDI-TOF MS and representative spectra are shown. (B) Graphical representation of Aβs generated over a temperature gradient, analysed by MALDI-TOF MS. Mean; N = 3 independent experiments. (C) Aβ 37/38/40/42 peptides and (D) total secreted Aβ from HEK293T cells expressing APP_C99_ WT, FAD, or FAD+D23F/N27A/K28A mutations, analysed by multiplex MSD ELISA. In panel C, Aβ peptides are shown as % of the sum of all measured peptides. Mean ± SD; N ≥ 3 independent experiments. (D) One-way ANOVA followed by Dunnett’s post hoc test with comparison to WT was used to determine statistical significance (P < 0.05). ****P < 0.0001, ***P < 0.001, **P < 0.01 F (DFn, DFd): F (6, 30) = 11.69. Unpaired t-tests were used to determine statistical significance between two specific conditions; (P < 0.05). ^#^ ^#^P < 0.01, ^#^P < 0.1, ns; not significant. Data was normalized to APP_C99_ expression levels determined by western blot (**Figure EV4C**). (E) Aβ38/42 ratio of WT and mutant APP_C99_ substrates from (C) normalized to WT. Unpaired t-tests were performed to determine statistical significance (P < 0.05). ****P < 0.0001, ***P < 0.001. ND = not determined since mutant substrates harbouring the K28A mutation lowered Aβ42 levels below detection level. (F) Secreted Aβ 37/38/40/42 analysis of WT or PSEN1 FAD MEF cell lines transiently overexpressing WT or mutant APP_C99_. Aβ peptides are shown as % of the sum of all measured peptides. The same colour code applies as in (C). Mean ± SD; N ≥ 3 independent experiments. (G) Aβ(37+38/40+42) ratio of (F). Unpaired t-tests were performed to determine statistical significance in comparison to the WT APP_C99_ in each respective MEF cell line (P < 0.05). ****P < 0.0001, ***P < 0.001. AAOs of FAD mutations listed in (**Table S1**). Mutations marked with an asterisk (*) in panels C and F drastically lowered Aβ generation, but Aβ profiles could still be determined.

The analysis also demonstrated generation of shorter Aβ peptides from both mutant substrates (K28A, K28F) at all tested temperatures (**Fig 4B**), compared to the WT, indicating that the more efficient processing of these substrates steams from mutation-driven stabilization of E-S interactions. We additionally tested the D23V+K28V and N27V+K28V APP_C99_ mutants in the detergent-based assay (**Fig EV4B**), due to their strong effects on GSEC processivity in cells (**Fig 2F**). Both mutants rescued the destabilizing effects of detergent to different degrees; with the D23V+K28V mutant producing Aβ peptides as short as Aβ29 (**Fig EV4B**, upper panels). This demonstrated a synergistic effect of residues 28 and 23 in APP_C99_-ECD on E-S stability and GSEC processivity.

### APP_C99_-ECD destabilizes GSEC-Aβ interactions, and mitigation of this detrimental feature rescues the effects of pathogenic mutations on APP processing

We next investigated whether mutations in APP_C99_-ECD that enhance GSEC processivity rescue alterations in Aβ processing caused by AD-linked variants in APP or PSEN that impair E-S stability (Szaruga *et al*, 2017). We selected pathogenic mutation for which previous analyses demonstrated that their significant destabilizing effects shift Aβ profiles: APP-T43I, APP-I45F, PSEN1-L166P, PSEN1-G384A and PSEN1-Ins113T (also known as Intron 4) (Szaruga *et al*, 2017) (**Appendix Table S1**). In addition, PSEN1 mutations were chosen to map to different locations within PSEN. To test the effects exerted by APP_C99_-ECD, we selected the single K28A, D23F and N27A substitutions, as they modulate GSEC processivity to different degrees (**Figs 1D-G**).

Aβ profiles generated from FAD-linked APP -T43I and -I45F substrates showed enhanced generation of Aβ42 and Aβ38 peptides (**Fig 4C**), due to mutation-driven shifts towards the Aβ42 product line (Bolduc *et al*, 2016; Kumar-Singh *et al*, 2000; Chávez-Gutiérrez *et al*, 2012). Expression levels were robust for all substrates (**Fig EV4C**); nevertheless, we observed decreased total Aβ production for all FAD-linked substrates, relative to the WT (**Fig 4D**), as previously reported (Kumar-Singh *et al*, 2000; Guardia-Laguarta *et al*, 2010; Chávez-Gutiérrez *et al*, 2012).

The insertion of a second mutation (K28A, D23F or N27A) in APP_C99_-ECD promoted the conversion of Aβ42 into Aβ38, relative to the respective (single) FAD-linked mutant (**Fig 4C**), leading to significant increments in the Aβ38/Aβ42 ratio (**Fig 4E).** This particular ratio was not calculated for the K28A mutation since it drastically lowered Aβ42 production. However, its strong effects on Aβ processing were better assessed by the Aβ_total_/Aβ(37+38+40+42) ratio (**Fig EV4D)**. The K28A mutation also rescued the detrimental effects of the pathogenic (T43I and I45F) mutations on the global (endopeptidase) APP processing (K28A+T43I and K28A+I45F, respectively; **Fig 4D),** while the N27A substitution only rescued total Aβ levels generated from the T43I variant (N27A+T43I). These differences are likely explained by the milder effects of the N27A mutation, relative to the K28A (**Figs 1D-E and 4C**).

To investigate the processing of mutant K28A-, N27A-or D23F-APP_C99_ substrates by pathogenic L166P, G384A and Ins113T PSEN1 variants, we expressed WT or mutant APP_C99_ substrates in mouse embryonic fibroblasts (MEFs) expressing either WT or mutant GSEC complexes exclusively (Chávez-Gutiérrez *et al*, 2012; Szaruga *et al*, 2015). As reported, the tested PSEN1 variants increased Aβ42 production, while decreasing shorter Aβ(37/38/40) peptides, from the WT substrate (**Fig 4F)** (Chávez-Gutiérrez *et al*, 2012; Szaruga *et al*, 2015; Petit *et al*, 2022a). Processing of mutant (K28A, N27A, D23F) APP_C99_ substrates by the FAD-linked GSECs revealed enhanced production of shorter Aβ37/38 peptides, while lowering Aβ42 product levels (**Fig 4F)**, implying a higher degree of processivity relative to the WT. In support of this, significant increases in the Aβ(38+37)/(Aβ40+42) ratio and the Aβ_total_/Aβ (37+38+40+42) ratio demonstrated more efficient processing of mutant substrates in the FAD-linked cell lines, relative to the WT APP substrate (**Figs 4G and EV4E)**. In addition, total Aβ levels revealed significant increases in the global GSEC (endopeptidase) activity for the K28A mutant substrate (**Fig EV4F**). As seen before for the FAD-linked APP variants (**Fig 4D**), the milder N27 substitution did not rescue the effects of the pathogenic PSEN1 substitutions on the global GSEC activity levels (**Fig EV4F**). Collectively, these findings show that hydrophobic substitutions in APP_C99_-ECD (positions 23/27/28) counteract the destabilizing effects that pathogenic variants in APP (T43I and I45F) and PSEN (L166P, G384A and intron 4) exert on GSEC-Aβ interactions.

### DAPT and semagacestat act as competitive GSIs, and their paradoxical effects on Aβ profiles are facilitated by APP_C99_-ECD

Our findings led us to hypothesize that the destabilizing effects of APP_C99_-ECD on E-S interactions drive product dissociation and release. This notion is supported by molecular dynamics (MD) simulations analysing the stability of the GSEC-Aβ40 complex containing either WT or mutant Aβ40 peptides (Phe substitutions at positions Asp23, Asn27 and Lys28) showing that mutant complexes are energetically more stable than the WT E-S complex (**Fig EV4G**).

To challenge our hypothesis, we also took advantage of recent structural data mapping the binding pockets of ‘paradoxical’ GSIs (DAPT and semagacestat) within GSEC (Bai *et al*, 2015; Yang *et al*, 2021). These small compounds partially inhibit GSEC while causing paradoxical ‘FAD-mimicking’ increments in the production of longer Aβs (Qi-Takahara *et al*, 2005; Yagishita *et al*, 2006; Tagami *et al*, 2017), and recent cryo-EM structures revealed that both GSIs bind to the substrate-binding channel in PSEN (Bai *et al*, 2015; Yang *et al*, 2021) (**Fig 5A**). While semagacestat binds in close proximity to the active site (Yang *et al*, 2021), structural and modelling data suggest that DAPT binds more centrally in the channel (Aguayo-Ortiz *et al*, 2019; Bai *et al*, 2015). On these bases, we reasoned that DAPT and semagacestat could act as high-affinity competitive inhibitors to the substrates. A competitive mechanism, which is never 100% efficient, would explain the partial inhibition of the global GSEC activity; while the ‘paradoxical’ increase in production of longer Aβs would result from a more effective competition (displacement and release) of shorter Aβ substrates, than longer ones, due to their differential affinities towards the enzyme. Note that E-S stability/affinities progressively decrease during the sequential GSEC-mediated cleavage (Szaruga *et al*, 2017; Okochi *et al*, 2013).

**Figure 5.**
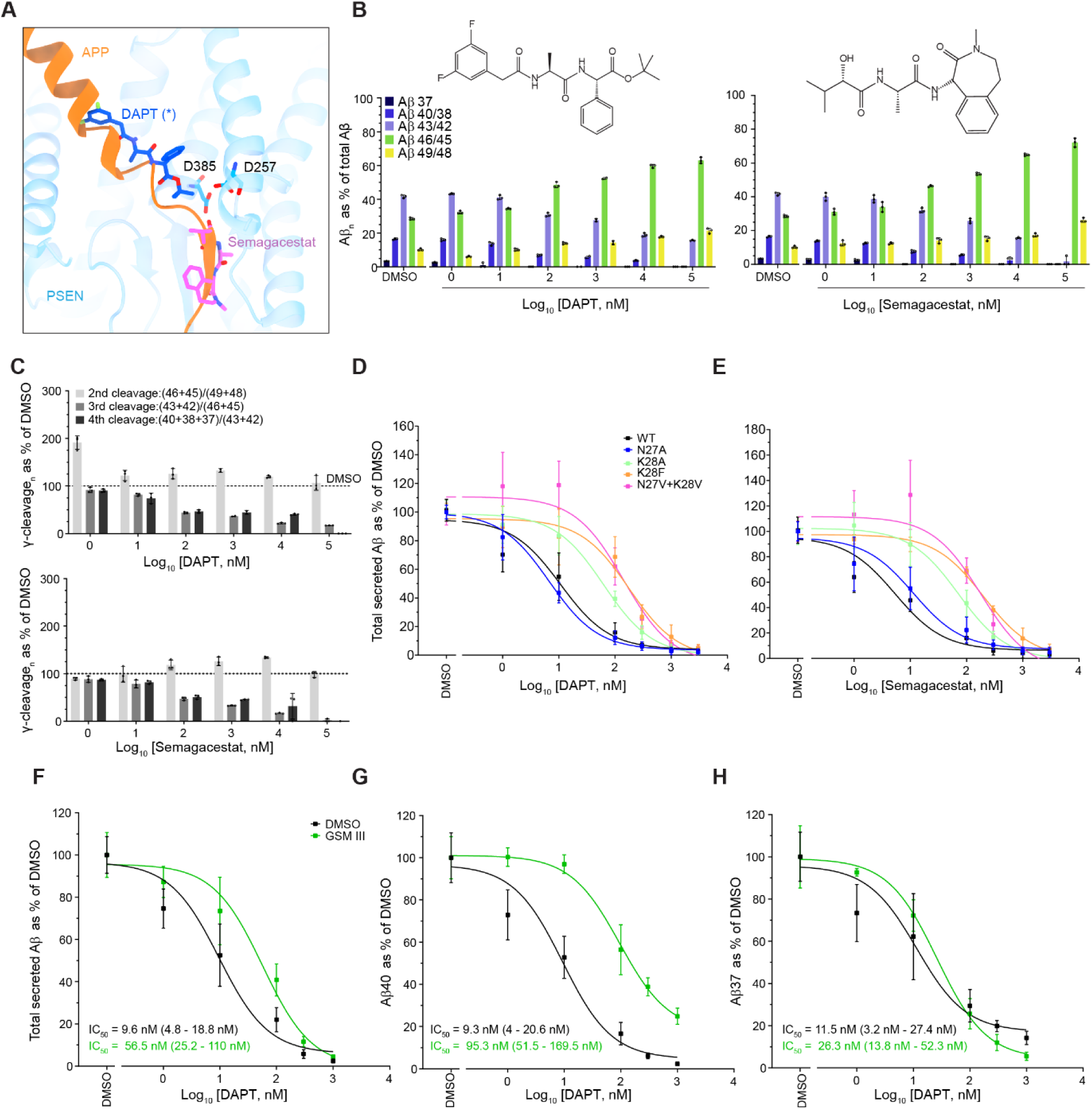
DAPT and semagacestat GSIs compete with substrates for binding to GSEC. (A) Superimposition of the GSEC-APP_C83_ (PDB: 6IYC), GSEC-semagacestat (PDB: 6LR4) and GSEC-DAPT (PDB: 5FN2) co-structures show GSI binding to the substrate binding pocket. DAPT (*) was not annotated in the high-resolution map (Bai *et al*, 2015) but electron densities and simulation data (Aguayo-Ortiz *et al*, 2019) support the shown binding pose. (B) Cell-free GSEC assays demonstrate that DAPT (left) and semagacestat (right) cause a relative increase of long Aβ species (Aβ ≥ 45), even at micromolar concentrations. Peptide product analysis was conducted by MALDI-TOF MS. Mean ± SD; N = 3 individual experiments. DAPT and semagacestat chemical structures are shown. (C) The efficiency of sequential γ-cleavage, assessed by the substrate/product Aβ peptide ratios from (B) reveals that the number of γ-cuts declines with increasing GSI concentrations, leading to (relative) enhanced production of longer Aβs (≥ 45). (D) Total secreted Aβ peptides in CM of HEK293T cells expressing WT or mutant APP_C99_ and treated with increasing concentrations of DAPT or (E) semagacestat. Mean ± SD; N ≥ 3 individual experiments. (F/G/H) Total secreted Aβ, Aβ40 and Aβ37 peptide levels in CM of HEK293T cells expressing WT or mutant APP_C99_ treated with either vehicle (DMSO) or 10 µM of GSM III, and increasing concentrations of DAPT. Mean ± SD; N ≥ 3 independent experiments.

Following this reasoning, we investigated if mitigation of the product release promoting effects of the APP_C99_-ECD would result in more stable E-S complexes and thereby hamper DAPT and semagacestat mediated-inhibition. We measured the inhibitory effects of these compounds in well-controlled *in vitro* assays using purified GSEC and APP_C99_. As previously reported (Yagishita *et al*, 2006; Qi-Takahara *et al*, 2005; Tagami *et al*, 2017), DAPT and semagacestat partially inhibited APP processing, while leading to increments in the production of longer Aβ species (**Figs 5B and EV5A-EV5C**). Moreover, dose-dependent effects (mainly) resulted in relative increases in Aβ46/45 production (**Figs 5B-C and EV5A**), which is in line with previous studies (Qi-Takahara *et al*, 2005; Yagishita *et al*, 2006; Tagami *et al*, 2017). In contrast, the transition-state analogue (TSA) inhibitor InhX efficiently inhibited all GSEC-mediated cleavages, as indicated by Aβ profiles (**Figs EV5A-C**).

We next assessed the effects of DAPT and semagacestat on the processing of the N27A-, K28A-, K28F-, and N27V+K28V mutant substrates by assessing total secreted Aβ levels in CM of HEK293T cells overexpressing these substrates. We selected mutations in APP based on their differential effects on processivity. The inhibitory profiles (**Figs 5D-E, respectively**), and derived IC_50_ values (**Appendix Table S2-S3**) demonstrated significant changes in the inhibitory potencies of DAPT and semagacestat for all tested mutant substrates, besides N27A, and relative to the WT condition. As mentioned above, the milder effects of the N27A substitution (**Fig 1D-E, 2D-E and 4C-G)** probably explain the lack of significant effects. Importantly, IC_50_ values determined for InhX treatment did not statistically differ from the WT IC_50_ (**Fig EV5D and Appendix Table S4**). The observed shifts in GSI potency are consistent with a competitive model where binding of DAPT or semagacestat to the substrate-binding channel, either blocks the entry for transmembrane GSEC substrates or leads to displacement (and release) of partially digested peptides.

### GSM binding counteracts Aβ product release facilitated by APP_C99_-ECD

Previous analyses have shown that imidazole-based GSMs act as stabilizers of GSEC-Aβ interactions (Petit *et al*, 2022c; Okochi *et al*, 2013) and lower Aβ product dissociation (Okochi *et al*, 2013). Our recent analyses locate the binding pocket of a potent imidazole-based GSM (GSM III) at the E-S interface, between PSEN loop 1 and the herein investigated region in APP (Petit *et al*, 2022c). We thus asked whether binding of the hydrophobic GSM III to E-S complexes reduces GSI potency. HEK293T cells overexpressing WT APP_C99_ were treated with 10 µM GSM III and increasing concentrations of DAPT. As before, we determined DAPT inhibitory profiles from the analysis of secreted, total Aβs (**Fig 5F**). We found a significant shift in IC_50_ values in the presence of GSM III, relative to vehicle (DMSO: 9.6 nM; 95% CI: 4.8 - 18.8 nM vs. GSM III: 56.5 nM; 95% CI: 25.2 - 110 nM). Interestingly, the shift is mainly driven by changes in the inhibition of Aβ40 (DMSO: 9.3 nM; 95% CI: 4 to 20.6 vs. GSM III: 95.3 nM; 95% CI: 51.5 to 169.5) (**Fig 5G**), rather than Aβ37 or Aβ38 (**Figs 5H and EV5E, respectively and Appendix Table S5)**, indicating that GSMIII mainly stabilizes the GSEC-Aβ40 complex and promotes its conversion to Aβ37.

## Discussion

Despite a high pathophysiological relevance, the molecular underpinnings of the processive proteolysis of Aβ by GSECs remain unclear. While various mutations in GSEC or APP have been found to impair APP sequential processing, thus promoting the generation of longer and toxic Aβs, only a few cases enhance production of shorter Aβ peptides. A notable example is position 28 in APP_C99_-ECD, for which substitutions (Ala/Glu) have shown increases in the generation of very short peptides (≤ 37 aa) (Petit *et al*, 2019; Kukar *et al*, 2011; Jung *et al*, 2014; Ren *et al*, 2007; Page *et al*, 2010; Ousson *et al*, 2013).

These observations support the notion that sequence and/or structural determinants in the substrate play a major role in modulating its sequential processing by GSEC.

Here a mutagenic screen in cells expressing WT or mutant APP_C99_ substrates identified the positions Glu22-Asp23 and Ser26-Lys28 in APP_C99_ as key determinants for the processive GSEC-mediated proteolysis (**Figs 1D-G**). The introduction of Phe at these positions markedly increased the generation of shorter Aβs, while Ala substitutions resulted in milder shifts towards shorter Aβ peptides. Notably, the D23F and K28F mutations abrogated the generation of ‘canonical’ Aβ (37/38/40/42) peptides by promoting their (further) processing into shorter peptides (≤ Aβ37). The generation of very short Aβs, ranging in length from 34 to 20 aa, implies that these substitutions not only promoted efficient sequential proteolysis of Aβ, but also extended substrate processing likely by promoting (mutant) APP_C99_-ECD threading through the transmembrane substrate-binding channel. We note that assessment of these very short Aβ peptides relied on mass spectrometry and, although this is a well-established method to determine Aβ profiles (Kukar *et al*, 2011; Jung *et al*, 2014; Wanngren *et al*, 2012), the different ionization and aggregation propensities of these peptides might affect their detection. The product profiles presented here thus represent estimates of relative proportions.

Mutagenic analysis of the key APP_C99_-28 position showed that, except for Arg, all other aa substitutions promote the generation of shorter Aβs (< 37 aa) (**Figs 2A-C**), as assessed by the total-to-canonical Aβ_total_/Aβ(37+38+40+42) ratio. The fact that all substitutions in position 28 in APP_C99_, exempt for the conservative K28R, promoted enzyme processivity points at a suboptimal role of APP_C99_-Lys28 (WT) in Aβ processing. Furthermore, this ratio also revealed a direct correlation between hydrophobicity at position APP_C99_-28 and GSEC processivity.

Previous reports have suggested that a substrate-membrane anchoring interactions involving Lys28 and negatively charged phospholipids limit (Aβ) substrate accessibility to the active site (Jung *et al*, 2014; Kukar *et al*, 2011; Chen & Zacharias, 2022). While such an interaction may be at play, the drastic effects of the D23F mutation, and milder but consistent effects of hydrophobic substitutions at positions APP_C99_ -22 -26 and -27, on Aβ processing implicate the ECD of APP_C99_, besides Lys28, in the regulation of both the efficiency and extent of the GSEC-mediated sequential cleavage.

Given the proximity of Lys28 to the TMD of APP, we evaluated whether the effects of mutations may be mediated by an extension of the helical TMD by introducing single-, double-and tetra-Ala or Val key substitutions in APP_C99_-ECD. The analysis of Aβ profiles generated from Val mutant substrates (N27V+K28V, D23V+K28V, 4xV (25-28), 3xV (K16+D23+K28)) pointed at hydrophobicity in APP_C99_-ECD, rather than α-helical propensity, as the key determinant of the efficiency and extent of sequential GSEC-mediated Aβ proteolysis (**Figs 2D-F and EV2D-E**). In addition, Aβ profiles generated from substrates carrying more than one substitution showed a degree of cooperativity between the tested positions in APP_C99_-ECD. Structural analysis of amyloid fibres derived from human cortical tissue have shown an ionic interaction between Asp23 and Lys28 in Aβ40 (Ghosh *et al*, 2021). A salt bridge between these positions, however, does not seem to be relevant for Aβ processing since almost identical Aβ profiles are generated from the FAD-linked APP-D23N (Iowa) and WT substrates (Hunter & Brayne, 2018; Tomidokoro *et al*, 2010). Noteworthy, mutations at positions 23 and adjacent residues in APP_C99_ (e.g. 21 and 22) are causal for AD. The effect of these pathogenic variants is mostly linked to increased Aβ production (A21G or Δ19-24) (Haass *et al*, 1994; De la Vega *et al*, 2021; Tian *et al*, 2010), and/or altered aggregation of mutant Aβ peptides (E22G, E22Q) (Yang *et al*, 2018). Whether mutation-driven alterations in Aβ profiles occur and contribute to AD pathogenesis requires further investigation.

Intriguingly, our studies also implicate APP_C99_-ECD in the modulation of the global (endopeptidase) activity levels. This is best exemplified by the significant ∼3-fold increase in the total Aβ pool generated from the 4xV(25-28) mutant APP_C99_ substrate (**Fig 2D**). Total Aβ amounts can be used as a proxy for the endopeptidase activity levels given that they equal total AICD.

We also evaluated whether similar rules govern the GSEC-mediated processing of Notch, which mediates essential signalling events in pathophysiology. Our analysis show that Notch-His17 position is the counterpart of Lys28 in APP_C99_. Nβ profile analysis revealed substantial extended processing of the H17F mutant substrate (**Figs 3D-E)**, but reduced sequential cleavage of the H17K Notch mutant, all relative to the WT. In fact, the single H17F substitution was sufficient to facilitate the cleavage of the Notch TMD in full, supporting the notion that hydrophobicity at the juxtamembrane region of the substrate ECDs controls GSEC processivity by restraining substrate threading and promoting product release.

In this regard, the ∼10 aa shift in Nβ product length generated from the H17F versus H17K substrates, and the fact that charged Glu9 and Glu6 residues are located 9-12 aa upstream of position Notch-17 (**Fig 3A**) led us to speculate that substrate threading facilitated by the H17F mutation might have been ‘halted’ once charged Glu9/Glu6 residues reach the vicinity of the hydrophobic pore.

In the case of APP_C99_, we propose that Lys28 critically contributes to an ‘energy barrier’ that disfavours further processing of Aβ40 (∼12 aa downstream) and Aβ42/38 peptides (14 aa/10 aa) in the two major production lines (**Figs 1B and 1G**). When the first barrier (Lys28) is overcome – likely a stochastic event – substrate threading continues and fuels sequential proteolysis until the next energy barrier is reached (Glu22-Asp23 and potentially other polar/charged residues).

The data indicate that the K28V/F mutation only lowers the first energy barrier, while the D23V/F reduces the first and second barriers. Even though the K28V/F mutation efficiently promotes further threading and proteolysis, the intact second barrier restricts the generation of peptides shorter than 28 aa. In contrast, the D23V/F is not as efficient as K28V in overcoming the first barrier (which explains the generation of the relatively longer Aβ40/38 peptides), but does further extend proteolysis to generate very short peptides. Interestingly, when Lys16 (likely the third barrier) is mutated together with Lys28 and Asp23 (K16V+D23V+K28V, **Fig 2F**) even shorter peptides are generated.

Our (thermoactivity) analysis demonstrate that hydrophobic substitutions of Lys28 (K28A/F) promote processivity by stabilizing GSEC-Aβ interactions (**Figs 4A-B**), which is consistent with the ‘energy barrier’ model of product release. Importantly, hydrophobic substitutions in APP_C99_-ECD (K28A, D23F and N27A) rescues the destabilization induced by detergent-solubilization or aggressive FAD-linked mutations in PSEN1 (L166P, G384A and Intron4) or APP (T43I, I45F) (**Figs 4C-G**), though to different degrees. Our studies support a model in which polar/charged residues in the juxtamembrane of the substrate (APP_C99_-Lys28 and H17 in Notch) serve as a ‘pivot’ mechanism wherein other polar interaction in the ECD, potentially involving the solvent, contribute to and collectively modulate product release by destabilizing the interaction of the substrate with GSEC.

In addition, the data highlighted rescuing effects of the stabilizing mutations in APP_C99_-ECD on the global activities of GSECs or APP bearing FAD-linked mutations (**Figs 4D, EV4C and EV4F**). These findings, together with the increases in total Aβ generation observed for mutations in the juxtamembrane region of APP_C99_ (**Figs 2A and 2D**), support the involvement of APP_C99_-ECD in the regulation of global GSEC activity. Further research is however needed to address whether these effects are mediated by higher E-S affinity and/or turnover.

We then investigated the pharmacological implications of this substrate ECD driven mechanism. Structural data show semagacestat and DAPT bound in the transmembrane catalytic pore in PSEN1 (Aguayo-Ortiz *et al*, 2019; Bai *et al*, 2015; Yang *et al*, 2021). We reasoned that a competitive mechanism would explain the reported paradoxical and ‘FAD-mimicking’ activities of these GSIs on Aβ production (Qi-Takahara *et al*, 2005; Yagishita *et al*, 2006; Tagami *et al*, 2017) (**Figs 5A-C**). In this model, mitigation of the product release promoting effects of APP_C99_-ECD would hamper DAPT and semagacestat mediated-inhibition. Dose-response inhibitory profiles of DAPT and semagacestat revealed that APP mutations stabilizing E-S interactions (K28A, K28F and N27V+K28V) significantly increase the IC_50_ values of these GSIs (**Figs 5D-E**), supporting a competitive mode of inhibition.

Interestingly, we observed that DAPT-induced inhibition of total Aβ was shifted towards higher GSI concentrations when cells were treated with a potent GSEC modulator (GSM III, **Figs 5F-G**). These data thus raise the possibility that binding of (hydrophobic) GSM III to the extracellular/luminal E-S interface (Petit *et al*, 2022c) lowers GSEC-Aβ_n_ dissociation (Okochi *et al*, 2013) by adding hydrophobicity in proximity to the key juxtamenbrane region. GSM binding may not only trigger allosteric changes in the E-S complex (Takeo *et al*, 2014; Petit *et al*, 2022c) but also displace water molecules from the E-S juxtamembrane region and this lowers the desolvation penalty for burying polar side chains of the substrate in the hydrophobic catalytic pore (substrate threading).

Our previous studies show that the shortening of the Aβ_n_ substrate progressively weakens (sequential) E-S assemblies (Szaruga *et al*, 2017); therefore, the substrate-driven product release mechanism is likely to play a more prominent role as the sequential GSEC cleavage progresses. One may ask if APP_C99_-ECD acts as ‘pulling force’ that will ultimately lead to (Aβ) product release, what drives the processive γ-cleavages? We speculate that the newly generated carboxy-terminus of Aβ could exert an ‘inward’ force extending the negatively charged C-terminus of the *de novo* Aβ substrate towards the cytosolic environment, away from the negatively charged active site. This inward pulling mechanism would not only extend the substrate and facilitate substrate fitting into S1’-S3’ pockets (Bolduc *et al*, 2016), but potentially contribute to the threading mechanism that fuels the processive GSEC cleavage (‘tug of war’ model).

In this regard, MD simulations with either WT Aβ40_coo-_ or a mutant presenting a neutral C-terminus Aβ40_NME_ in complex with GSEC showed that WT Aβ40 is energetically more favoured to transition from a product (P) to a cleavable substrate (S) position (ΔΔG_P→S_ = -13.9 kcal/mol), compared to a C-terminally neutral Aβ40 (ΔΔG G_P→S_ = -2.4 kcal/mol) **(Figs 6A-B**). We estimate that the difference in free energy for WT Aβ may be sufficient to break ∼7 hydrogen bonds (approximately 1.93 kcal/mol per hydrogen bond in a α-helix (Sheu *et al*, 2003)), and thus it is reasonable to consider a contribution to the threading of the substrate in a ‘tug of war’ model (presented in **Fig 6C**). Indeed, recent MD data supports the view that upon the first ε-cut a charged substrate carboxyl-terminus appears in Aβ49 (Bhattarai *et al*, 2022). Our MD simulations further indicate that interactions between the C-terminal COO^-^ moiety in Aβ and the positively charged Arg377 and Lys380 in PSEN1 contribute to the ‘extension’ mechanism (**Figs 6A and Appendix Figure S1 A-B**). PSEN1- Arg377 and Lys380 are conserved from invertebrates to humans (**Appendix Figure S2**), supporting their potential key involvement in the sequential proteolytic mechanism.

**Figure 6.**
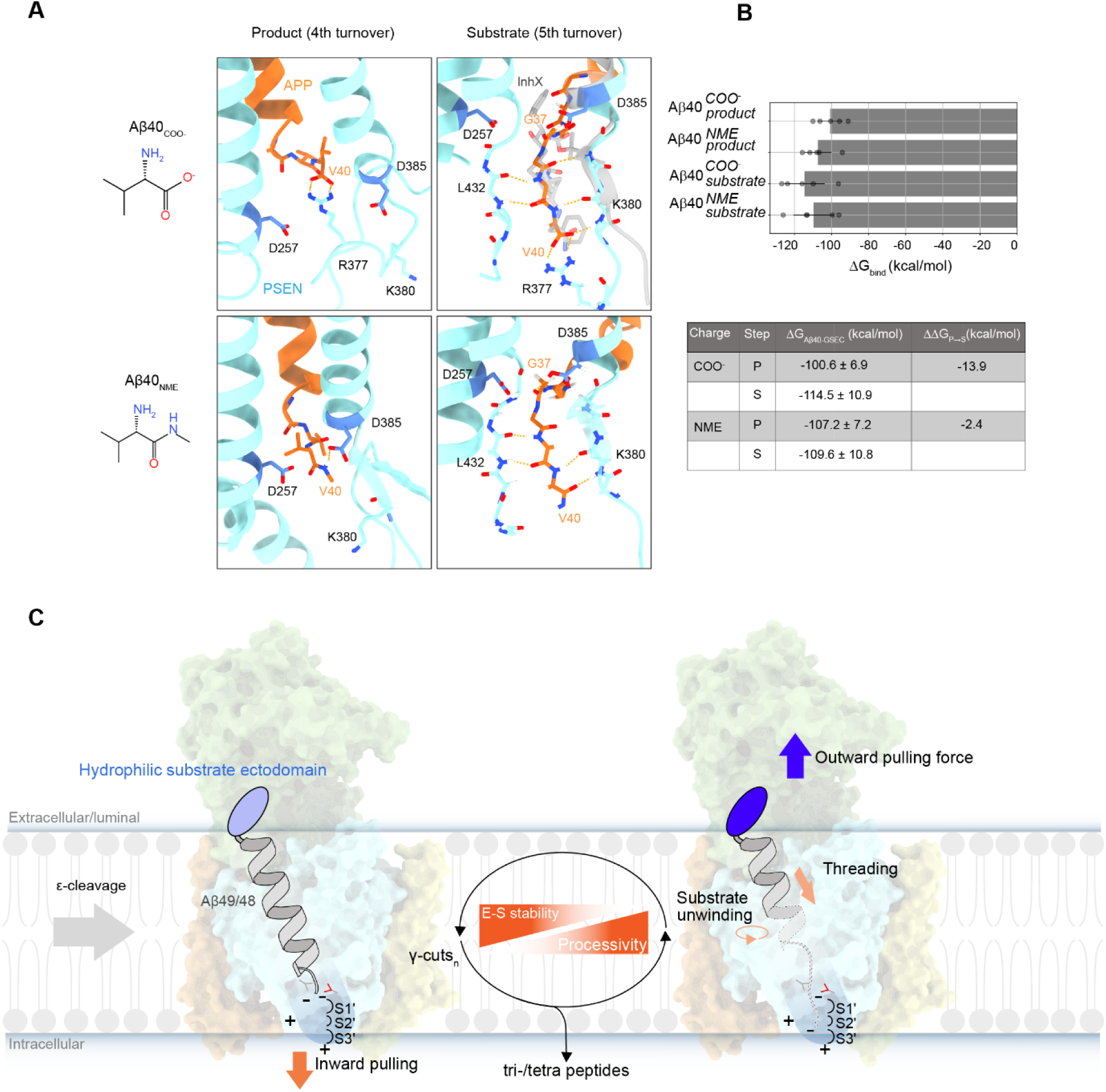
The ‘tug of war’ model of sequential GSEC proteolysis. (A) Representative MD snapshots of the GSEC-Aβ40 complex at the GSEC active site. Aβ40 as product (P, left) or substrate (S, right), with charged (COO) or neutral C-terminus (NME) shown in upper and lower panels, respectively. The data suggest that the charged C-terminus engages in interactions with PSEN1, specifically with the positively charged PSEN1-R377 (see upper panels), which facilitates product → substrate (P → S) transition. Structural data of the GSEC-InhX complex (PDB: 7C9I) show the TSA InhX (depicted in grey in right-upper panel) establishing similar interactions with PSEN1, supporting the MD simulations. (B) Binding free energies derived from MD simulations for GSEC with either WT (COO-) or neutral (NME) Aβ40 peptides interacting as products or substrates, respectively. ΔG_bind_ values indicate that (WT) Aβ40-COO^-^ is about six times more favoured to transition from P to S state, compared to neutral Aβ40-NME (ΔΔG_P→S_ = -13.9 kcal/mol vs. ΔΔG_P→S_ = -2.4 kcal/mol). N = 5, mean ± SD. (C) The ‘tug of war’ model of the GSEC-mediated processing. Polar interactions involving the ECD of the substrate destabilize E-S complexes. Since the stabilities of GSEC-APP/Aβ complexes progressively decrease during the sequential cleavage, the ’outward’ pulling force exerted by the ECD of the substrate becomes more apparent with the shortening of Aβ. On the intracellular side, **i**) the repulsion between the negatively charged C-terminus of Aβ substrates (generated *de novo* with each ε/γ-cut) and the negatively charged catalytic aspartate in PSEN1; and **ii**) the attraction between the negatively charged substrate terminus and positively charged residues present in PSEN1 (e.g. PSEN R377), collectively exert an ’inward’ force that pulls the C-terminus of Aβ towards the aqueous intracellular environment (blue gradient). Each γ-cut promotes (further) unwinding of the substrate TMD, and this ’inward’ pulling force facilitates its extension C-terminally and filling of the S1’-S3’ pockets. When the ‘energy barrier’ created by the polar substrate ECD is overcome (stochastic event) further substrate threading occurs and proteolysis is facilitated by the ’inward’ pulling force. The scheme in the middle illustrates the sequential nature of this process. When the (substrate-driven) ’outward’ pulling force overcomes E-S stabilizing interactions, occurring within the membrane-core, product release occurs.

In conclusion, the presented data assign a pivotal role to the substrate ECD in controlling both the efficiency and extent of the GSEC-mediated sequential processing. We show that the substrate ECD driven destabilization of E-S interactions underlies the product-release mechanism and has pharmacological implications as i) it facilitates the ‘paradoxical’ effects of some GSIs (such as DAPT and semagacestat) on Aβ processing and ii) its mitigation overcomes major E-S disrupting challenges linked to the effects of AD-causing mutations in *PSEN1* and *APP*. Our findings point at this mechanism as a sweet spot for the development of pharmacological strategies selectively targeting APP/Aβ processing in AD therapy.

## Materials and Methods

### Antibodies and compounds

ELISA capture antibodies (JRD/Aβ37/3 for Aβ37, JRF AB038 for Aβ38, JRF/cAb40/28 for Aβ40, JRF/cAb42/26 for Aβ42), Aβ N-terminal detection antibody (JRF/AbN/25) and imidazole-based GSM III modulator (synthesis described in (Velter *et al*, 2014)) were obtained through collaboration with Janssen Pharmaceutica NV (Beerse, Belgium). The anti-APP/Aβ 6E10 (#803003; epitope: 1-16), 4G8 (#800703; epitope: 17-24) and anti-HA. 11 epitope tag (#901514) antibodies were purchases from Biolegend. The monoclonal anti-FLAG antibody (#F3165) was ordered from Sigma-Aldrich. The 82E1 anti-human Aβ antibody (#10323) was purchased from Tecan. Complete protease inhibitor (PI) tablets (#11836145001), DAPT (#D5942) and semagacestat (LY450139; #SML1938) were purchased from Sigma-Aldrich. GSEC inhibitor L-685,458 was purchased from Calbiochem (#565771) and Bio-connect (#HY-19369). The polyethylenimine (linear, MW 25000, transfection grade (PEI 25K™) cell transfection reagent was purchased from tebu-bio (#23966-100).

### Generation of mutant APP and Notch substrates

Mammalian expression pSG5-APP_C99_-3xFLAG or pSG5-mouseNotch-3xFLAG constructs were subjected to site-directed mutagenesis using the QuikChange II XL Site-Directed Mutagenesis Kit (#200522) from agilent. Sequencing confirmed the introduction of the respective mutations.

### Determination of γ-secretase activity and processivity in HEK cells

HEK293T cells were plated at ∼70% cellular density. The next day, cultures were transiently transfected with pSG5-APP_C99_-3xFLAG constructs using a 1 mg/ml polyethylenimine (PEI) solution with a DNA:PEI ratio of 1:3. The medium was refreshed at one day post-transfection, cells placed back into the incubator for 24-30 h and collected. In case of GSM III or GSI (L-685,458 (InhX), DAPT or semagacestat) treatment, either compounds or vehicle (DMSO) were added to the media. The concentrations of Aβ peptides Aβ(37/38/40/42) in conditioned media were measured by a Meso Scale Discovery 4 spot ELISA (MSD ELISA). Alternatively, total secreted Aβ peptides were assessed by MSD ELISA using the 4G8 antibody (epitope: Aβ17-24) as capturing antibody (instead of the C-terminal specific anti-Aβ antibody).

### γ-Secretase in vitro thermoactivity assays

Proteolytic reactions were performed using purified ∼10 nM PSEN1/APH1B γ-secretase complexes and purified recombinant FLAG tagged substrates in 0.25% CHAPSO, 2.5% DMSO, 0.1% Phosphatidylcholine, 150mM NaCl and 25mM PIPES for 20 min. Enzyme mixes (containing all components except the substrate) and substrate dilutions were pre-incubated separately at the indicated temperatures for 5 min. In case of the thermoactivity assays a temperature gradient ranging from 37°C-58.5°C was applied. After pre-incubation, the substrate was added to the enzyme mix and proteolysis proceeded for 20 min. The APP intracellular domain product levels (AICD-3xFLAG) were analysed by SDS-PAGE and western immunoblot using the M2 anti-FLAG antibody, signals were quantified with an infrared imaging system. Data was normalized as % of WT conditions (WT APP_C99_ substrate). The final substrate concentrations in assays were saturating at 1.5 µM C99-3xFLAG. Additionally, *in vitro* activity assays, as described above, were carried out in presence of increasing concentrations of the indicated GSI (InhX, DAPT or semagacestat) and Aβ products assayed by MALDI-MS. Activity assay using detergent resistant membranes (DRMs) were performed as previously described (Szaruga *et al*, 2017) at 37°C and incubated for 90 min. DRMs were prepared from Hi5 insects cells, which overexpressed all four GSEC subunits.

### Detection of Aβ product profiles in Urea gels

Aβ products were analyzed in urea-based bicine/tris SDS–PAGE, as described previously (Wiltfang *et al*, 2002; Szaruga *et al*, 2017). Gel thickness was 0.75 mm and the composition of the separation gel was as follows: 8M Urea, 11%T / 5% C polyacrylamide, 0.4M H_2_SO_4_, 0.25% SDS, pH = 8.1. Electrophoresis was conducted at constant 100V for around 2h, after that, gels were transferred to a PVDF membrane and western immunoblot with 82E1 antibody, biotinylated anti-mouse IgG and streptavidin-HRP was performed. Signals were detected using ECL chemiluminescence with the Fujifilm LAS-3000 Imager.

### Expression and purification of GSEC complexes and substrates in Hi5 insect cells

Human GSEC, APP_C99_ and Notch constructs were purified as previously described (Szaruga *et al*, 2017). Human WT PSEN1, NCSTN-GFP, APH1B and PEN2 cDNAs were cloned into the pAcAB4 or pOET1 transfer vector (BD Biosciences). Co-transfection of the transfer vector (containing the heterologous cDNAs) and flashBacGoldTM DNA (Oxford Expression Technologies) in Sf9 cells allowed homologous recombination and production of baculoviruses bearing the four essential subunits of the GSEC complex. A PreScission Protease cleaving site (LeuGluValLeuPheGln/GlyPro) and GFP were cloned at the C-terminal site of NCSTN. Protease complexes were expressed in Hi5 insect cells. Infected Hi5 cells were collected at 72 hr post infection and lysed in 2% CHAPSO (Anatrace) buffer (25 mM Pipes pH 7.4, 300 mM NaCl, 5% Glycerol, 1X Protease inhibitors (PI). Affinity purification was carried out using a high affinity anti-GFP nanobody covalently coupled to agarose beads (NHS-activated beads, GE Healthcare) in a 3:1 ratio (mg:ml). PreScission protease cleavage between NCT and GFP eluted untagged g-secretase complexes (buffer composition: 25 mM Pipes pH 7.4, 150 mM NaCl, 0.5% CHAPSO, 5% Glycerol, 1 mM EDTA and 1mM DTT). Finally, removal of the GST-tagged PreScission protease by immunoaffinity pulldown using Glutathione Sepharose 4B (GE Healthcare) was performed and the purity of GSEC complexes was assessed by SDS-PAGE and Coomassie staining (InstantBlue Protein Stain, Expedeon).

In the case of APP_C99_-3xFLAG the purification was performed in the same way as GSEC purification with the difference that the transfer vector was either pAcAB4 or pOET1. APP_C99_ was solubilized in buffer containing 1% n-Dodecyl-β-D-Maltopyranoside (DDM) as detergent (Sol-Grade; Cliniscience), (Final APP_C99_ buffer composition: 25 mM Pipes pH 7.4, 150 mM NaCl, 0.03% DDM, 5% Glycerol).

### Expression and purification of GSEC substrates in from HEK293T cells

In order to validate the mouse Notch-3xFLAG substrate by mass spectrometry, purification was performed from HEK293T cells, by transiently transfecting the cells with WT or mutant (H17F, H17K, H17D, H17Q) constructs. Cells were treated with either the TSA L-685,458 or vehicle (DMSO), harvested and resuspended in 50 mM Tris-HCl pH 7.6, 150 mM NaCl, 1% Nonidet P-40, 1X PI and incubated on ice for 1 h. Membrane-solubilized protein fractions were obtained by ultracentrifugation at ∼100000 g for 1 h at 4°C. FLAG-tagged recombinant substrates were purified by immunoaffinity chromatography using the anti-FLAG M2-agarose beads (Sigma), according to the manufacturer’s protocols. All substrates were eluted in 100mM glycine HCl, pH 2.8, 0.03% DDM and immediately neutralized to pH 7 by the addition of 1 M Tris-HCl, pH 8.0.

### Generation of WT and FAD-linked MEF cell lines and evaluation of WT/mutant APP processing

dKO hPSEN1/2 MEF cells were rescued with WT or respective mutant hPS1 GSECs (L166P or G384A hPS1 FAD variants) as described previously (Petit *et al*, 2019). Cells stably expressing WT/mutant PSEN1s were selected. WT/mutant PSEN1 MEFs were electroporated with WT or mutant APP_C99_, Aβ peptides were quantified by ELISA. Raw values were used to calculate the Aβ profiles, where the addition of all canonical peptides is considered as 100%.

### Quantification of Aβ production by ELISA

Aβ37, Aβ38, Aβ40, Aβ42 product levels were quantified on Multi-Spot 96 well plates pre-coated with anti-Aβ37, Aβ38, Aβ40 and Aβ42 antibodies obtained from Janssen Pharmaceutica using multiplex MSD technology. For assessment of total Aβ Single-Spot 96 well plates (# L15XA-6, Multi-Array 96 well plate) were coated with the 4G8 anti-APP/Aβ antibody (epitope: 17-24; Aβ numbering). MSD plates were blocked with 150 ml/well 0.1% casein buffer for 2 hr at room temperature (600 rpm) and rinsed 5 x with 200 ml/well washing buffer (PBS + 0.05% Tween-20). 30 µL of SULFO-TAG JRF/AbN/25 or 6E10 detection antibody diluted in blocking buffer was mixed with 30 µL of standards (Aβ37, Aβ38, Aβ40 and Aβ42 peptides) or reaction samples diluted in blocking buffer and 50 µL per well were loaded. After overnight incubation at 4°C plates were rinsed with washing buffer and 150 µl/well of MSD Read Buffer T (tris-based buffer containing tripropylamine, purchased from Meso Scale Discovery) was added. Plates were read immediately on MSD Sector Imager 6000.

### Immunoprecipitation of Aβ peptides from conditioned media

HEK293T cells cultured in 10 cm^2^ dishes in DMEM/F-12 medium supplemented with 10% FCS were transfected with wild type or mutant APP_C99_ expressing constructs. At day one post transfection, the cell medium was replaced with low-serum medium (DMEM/F-12 medium containing 2% FBS) and depending on the experiment either DMSO or L-685,458 at a final concentration of 2,5 µM was additionally added to the medium. 24–30 h after medium replacement conditioned media were collected. To improve MS peak intensity for hydrophobic Aβ peptides Tween-20 at a final concentration of 0.025% was added to conditioned media as previously described (Portelius *et al*, 2007). Aβ peptides were immunoprecipitated using the 4G8, 6E10, anti-HA or 82E1 antibody (4 μg/10 ml of conditioned media) overnight at 4°C on rotation. Then 40 μl Protein G agarose beads (pre-blocked in PBS/0.5% BSA/0.05% Tween 20 pH 7.4) were added and incubation continued for 3 h. Beads were washed in PBS/0.05% Tween 20 pH 7.4 two times. The last wash in PBS/0.01% Tween 20 pH 7.4 was performed to reduce the detergent concentration. Dry beads were frozen at −20°C and subsequently subjected to MS analysis.

### MALDI-TOF MS sample preparation and analysis Aβ peptides

Beads were resuspended in 15 μL of SA matrix solution (38 mg/mL in water/ACN/TFA 20/80/2.5 (v/v/v)), and 30 nM Aβ1-28 was added (internal standard, IS). The sample was vortexed for 1 min and centrifuged for 5 min at ∼1,000 g. The supernatant (matrix-analyte mix) was collected, and 1 μl (9 technical replicates) was applied on a MALDI AnchorChip Target (Bruker Daltonics, Billerica, MA, USA) using dried droplet preparation and air-dried. All mass spectra were acquired on a rapifleX MALDI-TOF mass spectrometer (Bruker Daltonics) equipped with a 10 kHz Smartbeam™ laser using the AutoXecute function of the FlexControl 4.2. In the case of detergent assays, reactions were mixed in a 1:1 ratio with the matrix solution, 30 nM of IS were added and then analysed as described above. The acquisition method was calibrated using a 1/1/1 (v/v/v) mix of protein calibration standard I, peptide standard II (both Bruker Daltonics), and Aβ calibration standard using quadratic calibration. Briefly, each spectrum was acquired in linear positive mode within the mass range of m/z 2,000 to 20,000 with a low mass gate at m/z 1,800. 25,000 laser shots were automatically accumulated for each sample by random walk. Mass spectrometer parameters were balanced for optimal resolution and sensitivity in the Aβ peptide mass range (4-5 kDa). Subsequently, mass spectra were Savitzky–Golay-smoothed and baseline-subtracted with Top-Hat method and internally single-point calibrated (using Aβ1-28-peak). Average MALDI-TOF MS profiles were generated from nine single spectra and peaks were detected form the resulting average spectra with a S/N > 3 using “SuperSmoother” method. All processing was done using R 4.0.4 (R Foundation for Statistical Computing, Vienna, Austria) and the MALDIquant package (v. 1.19.3, (Gibb & Strimmer, 2012)). Generally mass peaks within a mass error of 500 ppm (part per million) were annotated. For graphical summaries presented here, only peptide masses are shown which appeared in at least two independent experiments.

### Molecular dynamics simulations

Starting structures of the Aβ40_γ37_ (substrate state) for the MD simulations was taken from the homology models generated in our previous work (Chen & Zacharias, 2022). The starting structure of Aβ40_γ40_ (product state) was prepared by truncating the residues C-terminal to Val40 from the Aβ43γ40 from the homology models of our previous work (Chen & Zacharias, 2022). Phe mutations of Aβ40, including D23F, N27F, K28F, at the substrate state were introduced using tleap from AmberTools22 (Case *et al*, 2022) D385 is protonated and D257 is unprotonated in the Aβ43_γ40_ system, based on a recently published pH calculation (Guzmán-Ocampo *et al*, 2023). Both D257 and D385 are deprotonated in the Aβ40_γ40_ system repel the substrate C-terminus in the product state. The enzyme-substrate complexes were solvated with 400 POPC molecules and TIP3P water with 0.15M potassium chloride using OPM (Lomize *et al*, 2012) and CHARMM-GUI (Lee *et al*, 2016) servers. Interactions between atoms are described by ff14SB (Maier *et al*, 2015) for proteins, lipi21 (Dickson *et al*, 2022) for lipids, and TIP3P (Mark & Nilsson, 2001) for water molecules. Two variations of the substrate C-terminus in AmberTools22 (Case *et al*, 2022), including the charged carboxylic acid (COO-) and the neutral NME capping group, were used to investigate the influence of charge on substrate binding.

Each simulation underwent energy minimization and equilibration process before the production run. First, The energy of the simulation box was minimized with 10 kcal · mol^-1^ · Å^-2^ and 2.5 kcal · mol^-1^ · Å^-2^ positional restraint on the proteins and the lipid, respectively, for 7000 steps using pmemd from AMBER (Mark & Nilsson, 2001). Then, the simulation box was equilibrated for 400ps with gradually releasing positional restraint, from 10 kcal · mol^-1^ · Å^-2^ and 2.5 kcal · mol^-1^ · Å^-2^ to 2.5 kcal · mol^-1^ · Å^- 2^ and no restraint on protein and lipid, respectively. Five simulations of 200ns each were performed for data analysis. Integration time step was set to 4fs with the use of SHAKE algorithm (Andersen, 1983) and hydrogen mass repartitioning method (Hopkins *et al*, 2015) with a non-bonded cut-off distance of 9Å. A temperature of 303.15 K and a pressure of 1 bar were maintained using Langevin dynamics (Goga *et al*, 2012) and Berendsen barostate (Berendsen *et al*, 1984), respectively, in the equilibration and production run process using the GPU-accelerated pmemd package from AMBER (Salomon-Ferrer *et al*, 2013). The interaction free energy between Aβ40 and γ-secretase at the product state (Aβ40_γ40_) and at the substrate state (Aβ40_γ37_) were calculated with molecular mechanics coupled with generalized Born and surface area continuum solvation (MM/GBSA) method using MMPBSA.py (Miller *et al*, 2012). Only the last 100ns of each simulation was submitted for the energy calculation using GB model II (igb=5) (Onufriev *et al*, 2004). Because no lipid was present between Aβ40 and γ-secretase around the regions of interest, an external dielectric constant of 80 was used.

## Expanded view figures

**Figure EV1.**
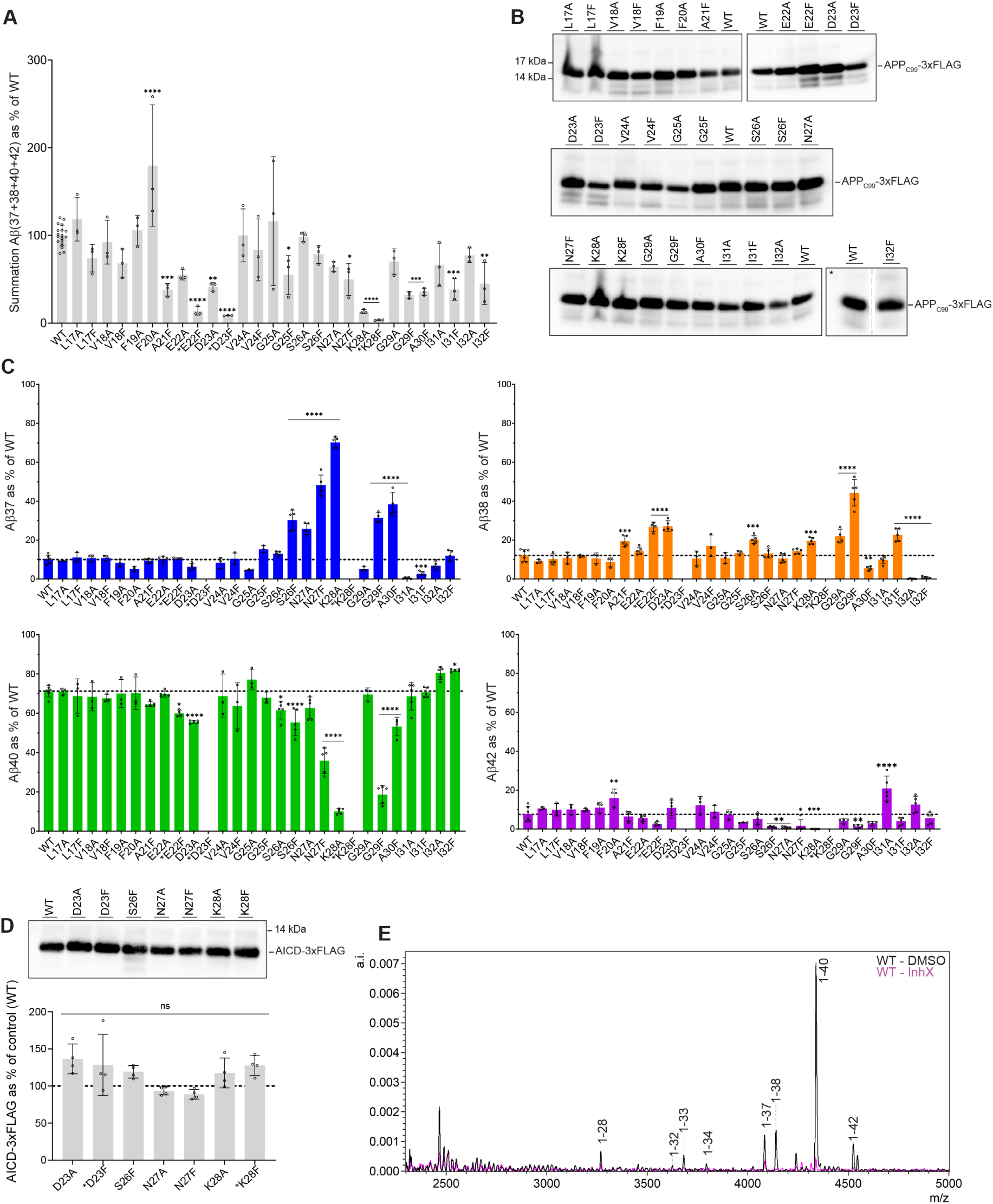
Effects of Ala/Phe APP_C99_ mutants on Aβ production and expression levels. (A) Summation of secreted Aβ(37+38+40+42) peptide amounts generated from HEK293T cells expressing WT or mutant APP_C99_ (from **Fig 1D)**, depicted as percentage normalized to WT. Mean ± SD; N = 3 independent experiments. One-way ANOVA followed by Dunnett’s post hoc test with comparison to WT was used to determine statistical significance (P < 0.05). ****P < 0.0001, ***P < 0.001, **P < 0.01 F (DFn, DFd): F (28, 73) = 10.06. (B) Representative western blot analysis of APP_C99_ 3x-FLAG levels in HEK293T cells expressing WT or mutant APP_C99_ substrates. Cell pellets were lysed with RIPA buffer and same volumes were loaded on SDS-PAGE followed up by western blotting with the anti-FLAG antibody and densitometric analysis.*Lanes in between WT and I32F were cropped out to condense the visualized blot (see bottom right WT and I32F) as indicated by the dashed line. (C) The proportion of secreted Aβ 37, 38, 40 or 42 peptides generated from WT or mutant APP_C99_ expressing cells (related to **Fig 1D)**. Mean ± SD; N ≥ 3 independent experiments. One-way ANOVA followed by Dunnett’s post hoc test with comparison to WT was used to determine statistical significance (P < 0.05). ****P < 0.0001, ***P < 0.001, **P < 0.01, *P < 0.1. (D) *De novo* generation of AICD-3xFLAG product levels was quantified in cell-free activity assays. Purified WT GSEC was incubated with saturating concentrations of purified WT or mutant (D23F, K28F) APP_C99_-3xFLAG substrates. As controls the corresponding Ala substitutions (K28A and D23A) and S26F, N27A or N27F APP_C99_ mutants were included. The upper panel shows a representative western blot analysis of AICD-3xFLAG. Quantifications are shown below. Mean ± SD; N = 4 independent experiments. One-way ANOVA followed by Dunnett’s post hoc test with comparison to WT was used to determine statistical significance (P < 0.05). ns; not significant F (DFn, DFd): F (7, 24) = 3.559. (E) Representative MALDI-TOF MS spectra of Aβs from CM of HEK293T cells expressing WT APP_C99_, IPed with the 6E10 antibody. Treatment with InhX abolishes Aβ generation (purple spectrum) compared to vehicle (DMSO) conditions (black spectrum).

**Figure EV2.**
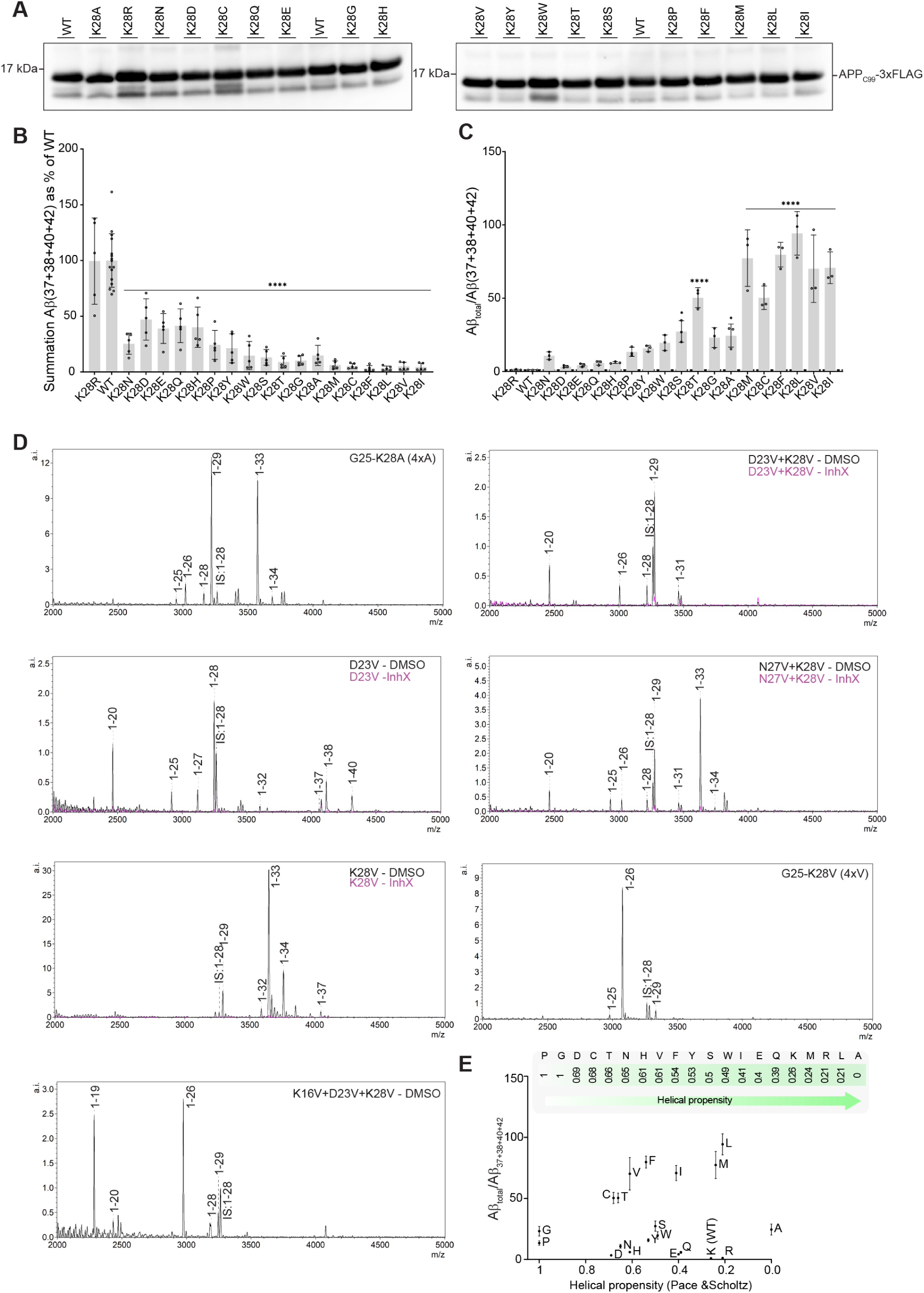
Mutagenesis of position 28 in APP_C99_-ECD. (A) Representative western blot of total lysates from HEK293T cells expressing WT/mutant APP_C99_ 3x-FLAG substrates using the M2 anti-Flag primary antibody. (B) Summation of Aβ(37+38+40+42) measured in CM of HEK293T cells transfected with WT or mutant APP_C99_ constructs and analysed by ELISA. Mean ± SD; N = 4 independent experiments. One-way ANOVA followed by Dunnett’s post hoc test with comparison to WT was used to determine statistical significance (P < 0.05). ****P < 0.0001 F (DFn, DFd): F (19. 91) = 28.45. (C) Aβ_total_/Aβ(37+38+40+42) ratios used in **Fig 2C**. Aβ_total_ and Aβ(37+38+40+42) peptides were quantified by 4G8 MSD ELISA and multiplex MSD ELISA, respectively. The WT ratio was set to 1, so that increments in shorter peptides (< 37aa) are > 1. Mean ± SD; N = 3 independent experiments. One-way ANOVA followed by Dunnett’s post hoc test with comparison to WT was used to determine statistical significance (P < 0.05). ****P < 0.0001, ***P < 0.001, **P < 0.01, *P < 0.1. F (DFn, DFd): F (19, 40) = 34.81. (D) Representative MALDI-TOF MS spectra of IPed Aβs from CM of HEK293T cells expressing WT or mutant APP_C99_ substrates (data related to **Fig 2F**). Cells expressing single and double valine mutant substrates were treated with InhX to confirm specificity of Aβ signals. (E) Correlation between WT/mutant Aβ_total_/Aβ(37+38+40+42) ratios and helical propensity of the amino acid substitution (Pace & Scholtz, 1998). AA are shown in one letter code.

**Figure EV3.**
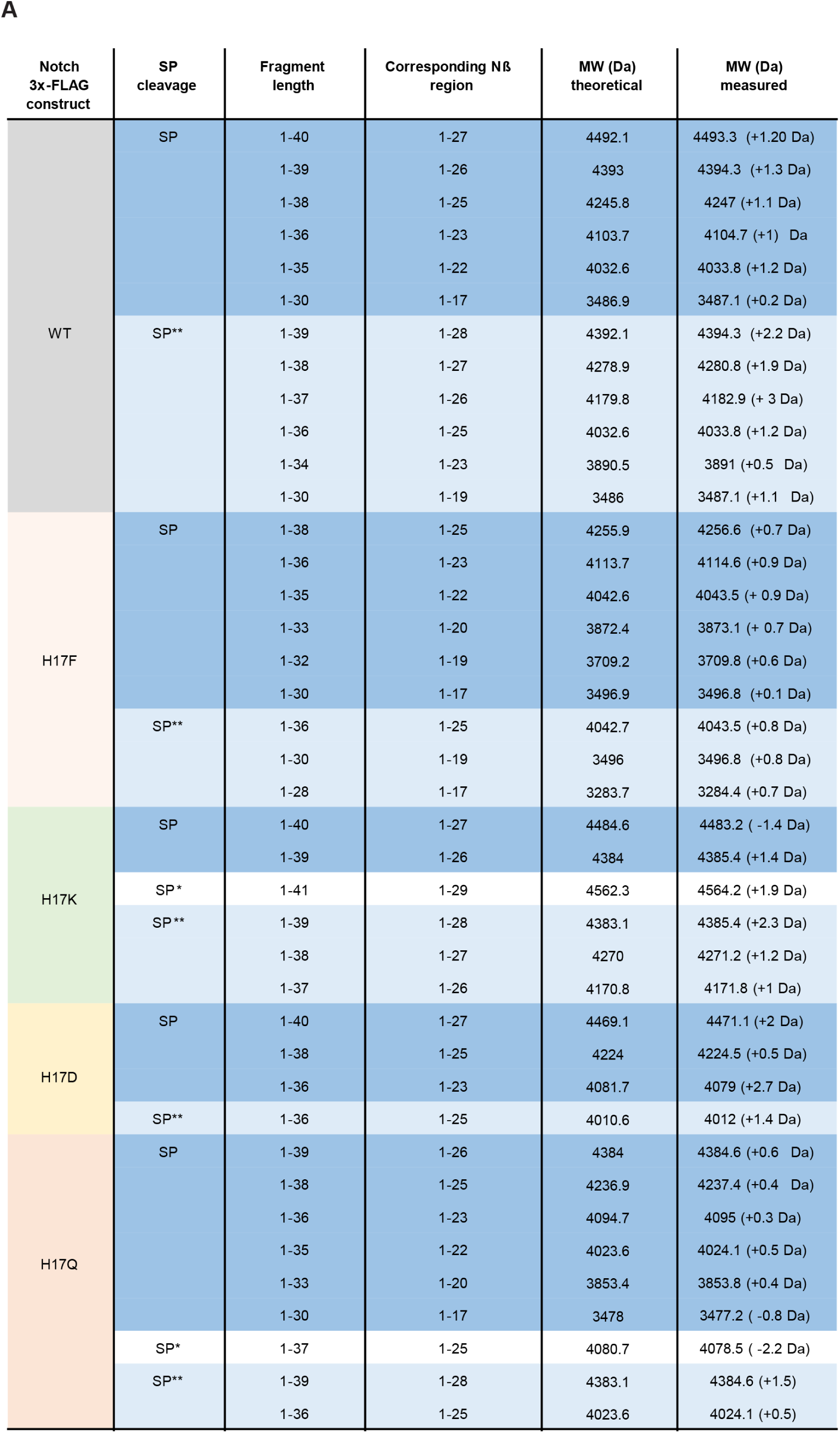
Mass spectrometry validation and analysis of mouse Notch. (A) Overview of detected masses and assigned Nβ fragments generated from WT and mutant Notch substrates. The respective fragment length and the corresponding length of the resulting Nβ peptide, when SP-and HA-tag associated residues are subtracted, are provided. Alternative SP cleavage leads to different Notch substrate lengths (see Fig 3); therefore, similar proteolytic cleavages result in distinct Nβ fragments masses. Only Nβ fragments are shown, of which a corresponding initial substrate mass (see **Fig 3C**) was detected.

**Figure EV4.**
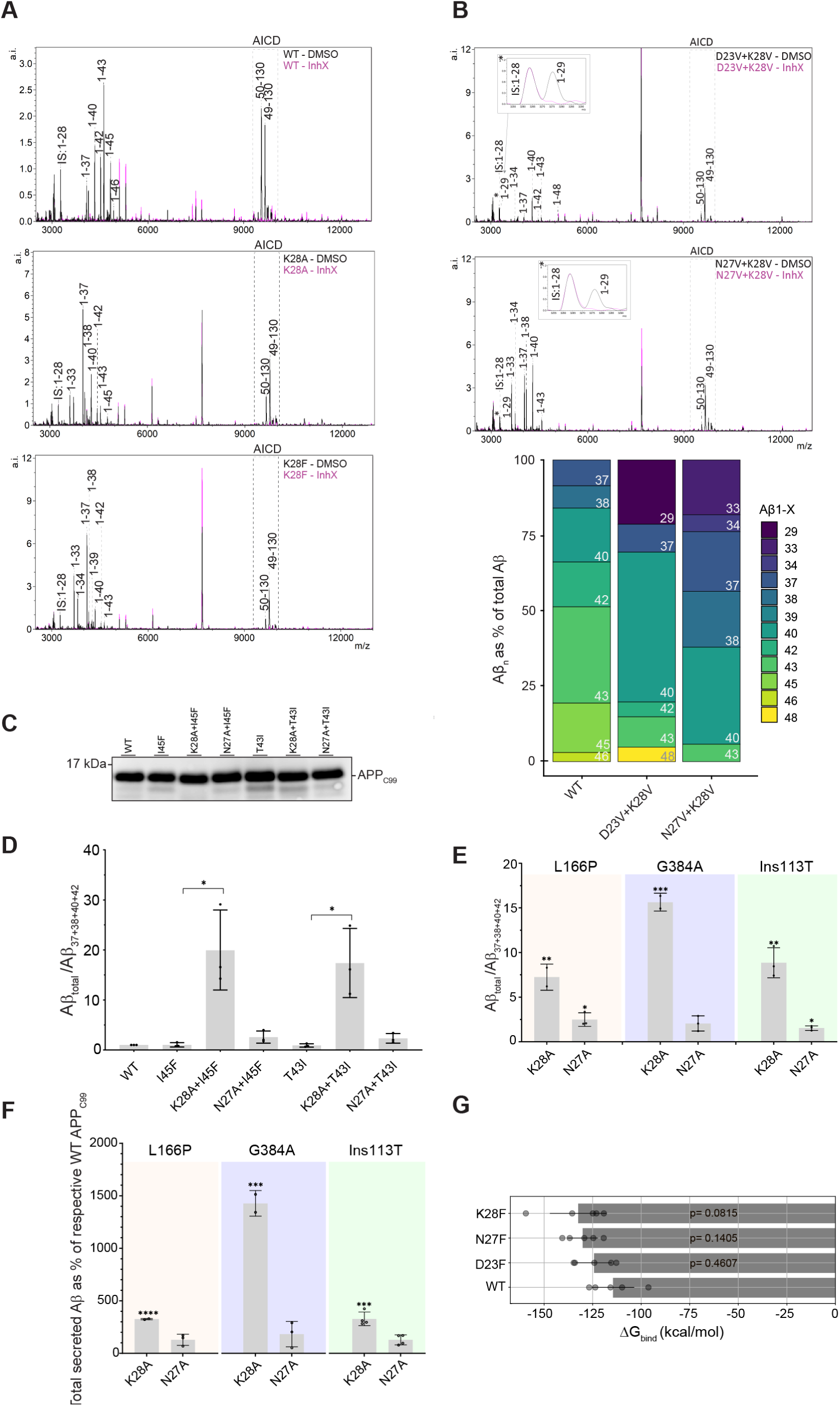
APP_C99_-ECD modulates sequential APP/Aβ processing and rescues effects of FAD mutations. (A-B) Representative MALDI-TOF MS spectra acquired by analysis of cell-free GSEC activity assays with either WT or mutant (K28A, K28F) APP_C99_. Assays were performed at 37°C in presence of DMSO vehicle or the GSEC inhibitor InhX (black and pink profiles, respectively). The lower panel in B shows a graphical summary of Aβ peptides measured in 4 independent experiments. Note that N27V+K28V produces Aβ1-29, however, due to a relatively low signal/noise (S/N) ratio it was not included in the graphical overview. (C) Representative western blot analysis of total lysates of HEK293T cells (from **Fig 4D**) expressing either WT or mutant APP_C99_ substrate. (D-E) Total secreted Aβ and secreted Aβ(37+38+40+42) peptides (Figs 4C, 4D, 4F **and EV4F**) measured by ELISA in CM of HEK293T or MEF cells expressing WT or mutant APP_C99_. The Aβ_total_/(Aβ37+38+40+42) ratio was calculated as an estimate of GSEC processivity. (D) Mean ± SD; N = 3 independent experiments. Unpaired t-tests; (P < 0.05). *P < 0.1. (E) Mean ± SD; N ≥ 2 independent experiments. Unpaired t-tests; (P < 0.05). ***P < 0.001,**P < 0.01, *P < 0.1. (F) Total secreted Aβ peptides from samples from **Fig 4F** were analysed using 4G8 ELISA. The APP_C99_-K28A mutation rescues impairments in total Aβ levels generated from the WT substrate by FAD L166P, G384A and Ins113T PSEN1/GSEC variants. Mean ± SD; N ≥ 2 independent experiments. Unpaired t-tests were used to determine statistical significance between two specific conditions. (P < 0.05). ****P < 0.0001, ***P < 0.001. (G) Free binding energies (ΔG_bind_) determined by MD simulations, between GSEC with either WT or mutant (D23F, N27F or K28F) Aβ40. Calculations were run in N = 5 independent attempts; Mean ± SD is shown. One-way ANOVA followed by Dunnett’s post hoc test with comparison to WT was used to determine statistical significance (P < 0.05). (DFn, DFd): F (3, 16) = 2.173. P-values are shown.

**Figure EV5.**
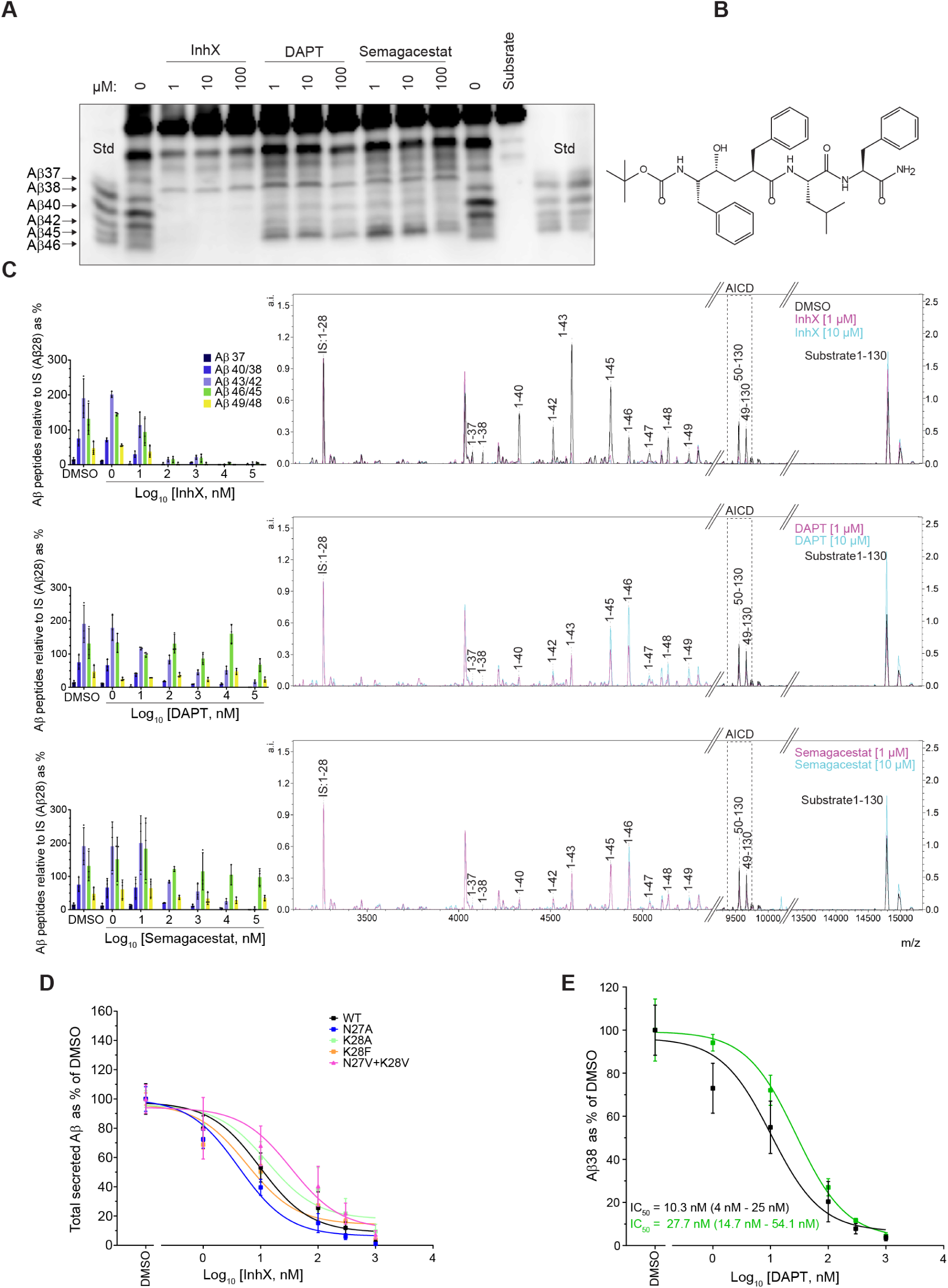
Competitive-like GSI action on GSEC in detergent and membrane conditions. (A) Representative urea gel/western blot analysis of cell-free GSEC activity assays performed with purified WT APP_C99_ and detergent resistant membranes (DRMs) derived from Hi5 insect cells expressing WT human GSEC (PSEN1/APH1B). Assays were incubated for 90 min at 37°C, InhX, DAPT or semagacestat were added at the indicated concentrations; as control vehicle (DMSO) was added. Purified APP_C99_ substrate or recombinant Aβ peptides at equimolar concentrations were loaded as (background) control or standards, respectively. Aβ profiles resolved in urea gels show enhanced generation longer species Aβ(45/46) in presence of DAPT and semagacestat, relative to DMSO. TSA InhX abolished production of all Aβ species. (B) Chemical structure of the GSI InhX. (C) GSEC activity assays using purified substrate and enzyme were performed in presence of vehicle (DMSO, black spectra) or GSIs (InhX, DAPT or semagacestat) at 1 µM or 10 µM (purple and blue, respectively). Representative MALDI-TOF MS spectra used in Aβ, AICD and substrate analysis is presented. On the left of each spectrum a summary of all detected Aβ peptides for each respective compound is shown, normalized to the internal standard (IS). Mean ± SD; N = 3 individual experiments (D) Total secreted Aβ peptides measured by ELISA in CM of HEK293T cells expressing WT or mutant APP_C99_ and treated with increasing concentrations of Inhx. Mean ± SD; N ≥ 3 individual experiments. (E) WT APP_C99_ was expressed in HEK293T, which were treated with either vehicle (DMSO) or 10 µM of GSM III, the Aβ38 peptide was quantified by MSD ELISA. Mean ± SD; N = 3 independent experiments.

## Appendix

**Appendix Figure S1.**
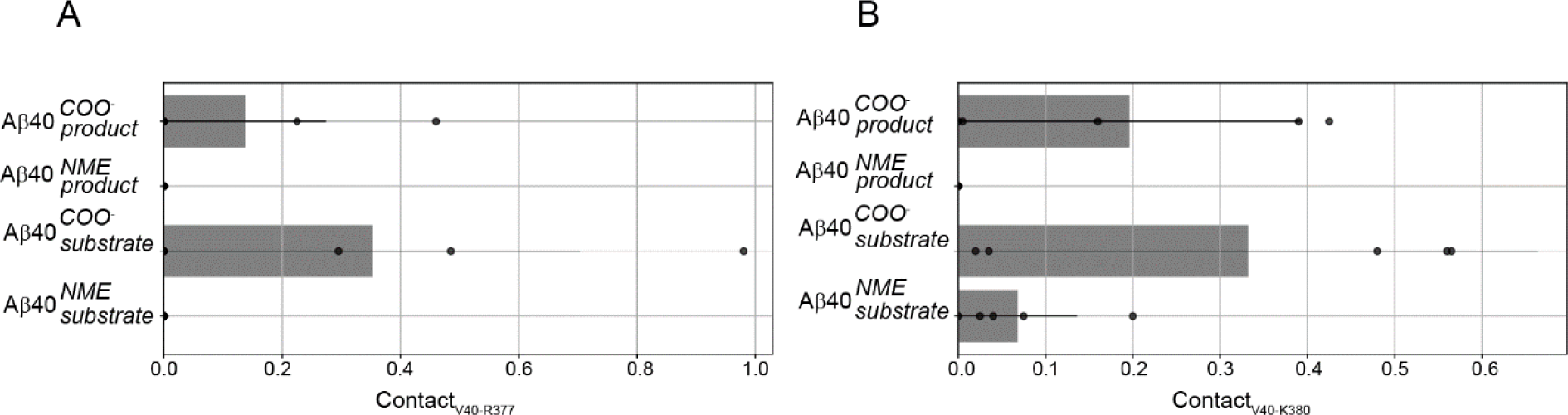
MD simulations reveal direct interaction between Aβ40 COO-and PSEN-R377/K380. (A) Fraction of contact between Aβ40-V40 and PSEN-R377 and (B) PSEN-K380 at the Aβ40 product and substrate state, respectively. Residues are considered in contact if their minimum intermolecular distance is ≤ 5 Å. Calculations were run in N = 5 independent attempts; Mean ± SD is shown.

**Appendix Figure S2.**
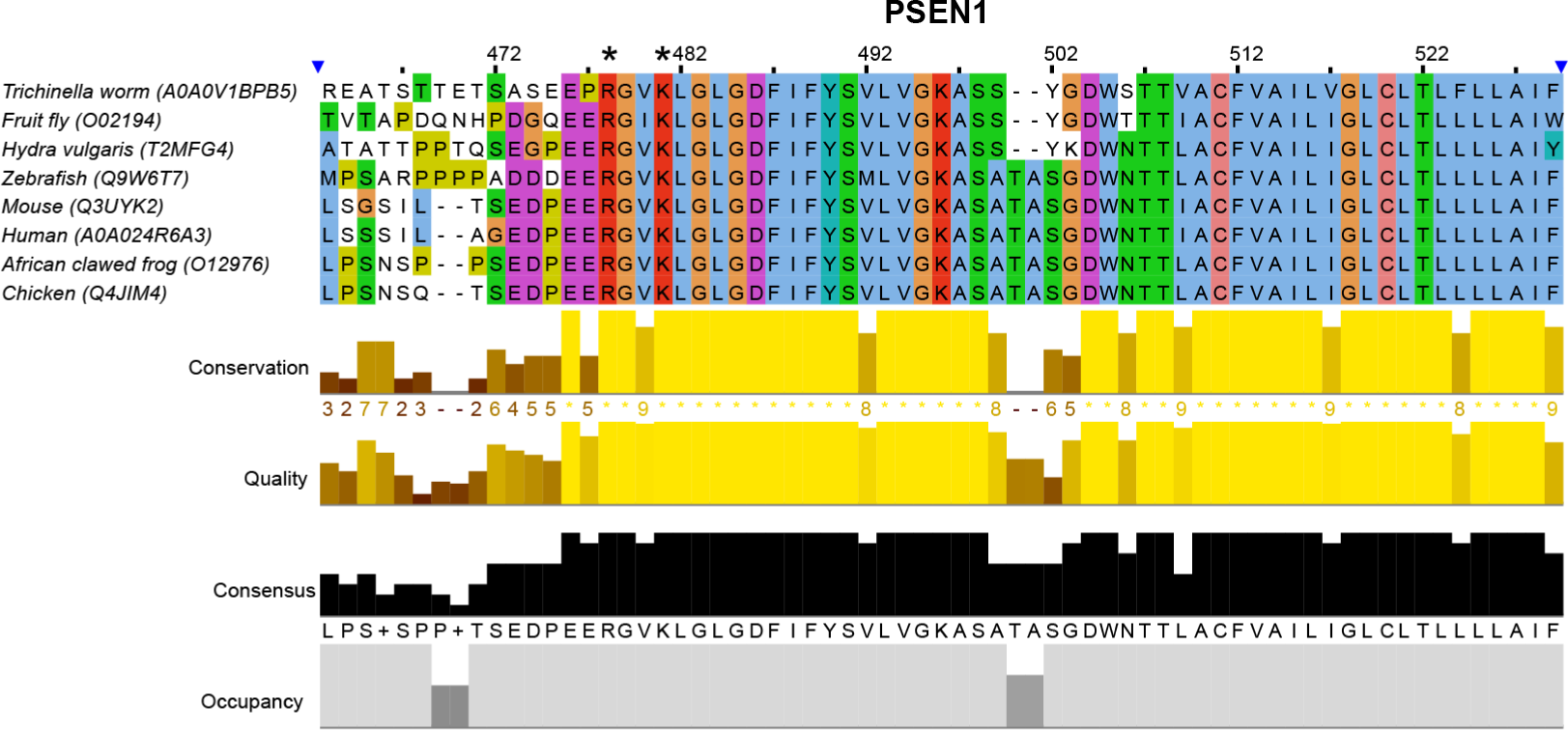
Multiple sequence alignment of PSEN1. Multiple sequence alignment of PSEN1 from different species using the Clustal Omega tool provided by EMBL-EBI (Madeira *et al*, 2022). The columns marked with an asterisk indicate the position of residue R377 and K380, respectively. Across the different species both positions are strongly conserved. The colours used follow the clustal color code (blue = hydrophobic; red = basic; magenta = acidic; green = polar; pink = cysteines; orange = glycines; yellow = prolines; cyan = aromatic; white = unconserved). For depiction of the alignment the jalview software was used (Waterhouse *et al*, 2009).

**Appendix Table S1.** FAD-linked PSEN1 and APP mutations utilized here. Table adapted from (Petit *et al*, 2022b).

| Protein | Mutation | Position | Mean AAO (range) |
| --- | --- | --- | --- |
| PSEN | L166P | TMD 3 | 23.5 (23-24) |
|  | G384A | TMD7 | 36.0 (26-45) |
|  | E280A | IC loop | 48.1 (46-52) |
|  | Ins113T | Loop 1 | 42.1 (35-45) |
| APP | T43I (Austrian) | APP-TMD | 34 (30-44) (Kumar-Singh *et al*, 2000; Edwards-Lee *et al*, 2005) |
|  | I45F (Iberian) | APP-TMD | 31 (Guerreiro *et al*, 2010) |

**Appendix Table S2.** Kinetic data of HEK293T cells with different concentrations of DAPT.

| APP<sub>C99</sub> substrate | IC<sub>50</sub> [nM] | IC<sub>50</sub> 95% CI [nM] |
| --- | --- | --- |
| WT | 10.7 | 6.4 to 17.5 |
| N27A | 6.7 | 5 to 9.1 |
| K28A | 63.7 | 46.1 to 83.6 |
| K28F | 177.4 | 93.6 to 324.3 |
| N27V+K28V | 142.2 | 88 to 227.2 |

**Appendix Table S3.** Kinetic data of HEK293T cell assays with different concentrations of Semagacestat or vehicle (DMSO)

| APP<sub>C99</sub> substrate | IC<sub>50</sub> [nM] | IC<sub>50</sub> 95% CI [nM] |
| --- | --- | --- |
| WT | 5.4 | 2.2 to 10.5 |
| N27A | 11.4 | 6.2 to 21 |
| K28A | 72.2 | 46.4 to 108.5 |
| K28F | 200.1 | 151.0 to 262.7 |
| N27V+K28V | 180.2 | 101 to 320.6 |

**Appendix Table S4.**
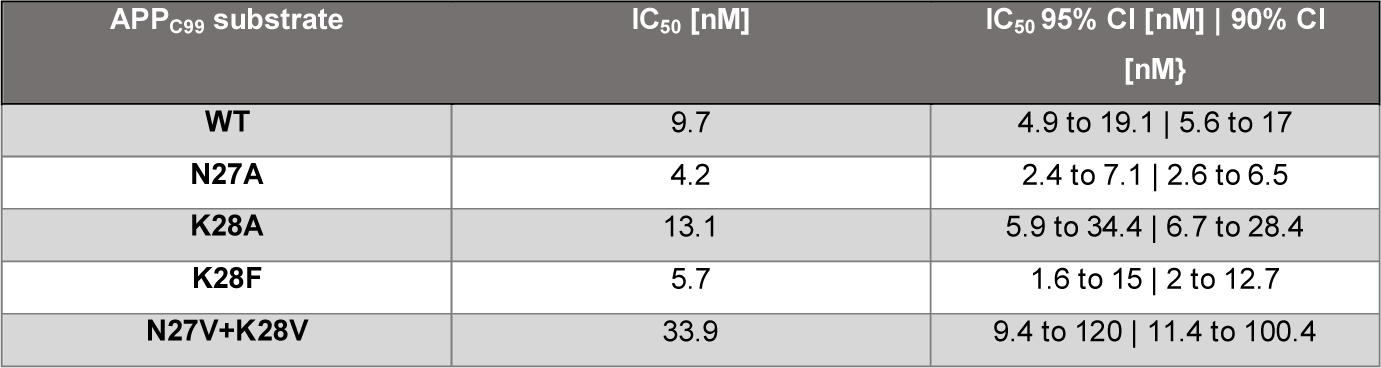
Kinetic data of HEK293T cells with different concentrations of InhX.

**Appendix Table S5.** Kinetic data of HEK293T cell assays with different concentration of DAPT or vehicle (DMSO) in presence or absence of GSM III.

| Condition | Aβ peptide measured | IC<sub>50</sub> [nM] | IC<sub>50</sub> 95% CI [nM] |
| --- | --- | --- | --- |
| DMSO | Aβ37 | 11.5 | 3.2 to 27.4 |
| GSM III | Aβ37 | 26.3 | 13.8 to 52.3 |
| DMSO | Aβ38 | 10.3 | 4 to 25 |
| GSM III | Aβ38 | 27.7 | 14.7 to 54.1 |
| DMSO | Aβ40 | 9.3 | 4 to 20.6 |
| GSM III | Aβ40 | 95.3 | 51.5 to 169.5 |
| DMSO | Total Aβ | 9.6 | 4.8 to 18.8 |
| GSM III | Total Aβ | 56.5 | 25.2 to 110 |

## Statistical analysis

Statistical analysis of the data was accomplished using the GraphPad Prism 9 software. Unpaired t-tests or one-way ANOVA with Dunnett’s post hoc test were used to test the significance of the changes as indicated in the figure legends. P-value < 0.05 was used as a predetermined threshold for statistical significance.

## Data availability

This study includes no data deposited in external repositories

## Acknowledgments

This work was funded by the FWO G0B2519N and G008023N research grants. C.H. acknowledges funding by BMBF (grant 13FH8I05IA (Drugs4Future)) within the FH-Impuls framework M^2^Aind. M.K. is supported by a PhD FWO fellowship (1S47020N). We thank Ivica Odorčić, Harald Steiner and Michel Vande Kerckhove for helpful advice and discussions. We also thank Lizzie Hill for critically revising the manuscript.

## Author contributions

L.C.G: conceptualization of the study, supervision of the experimental research, writing the manuscript. M.K: conceptualization of the study, performed experiments, analysed data, writing the manuscript. T.E and C.H.: performed mass spectrometry data collection, processing and interpretation. S.-Y.C and M.Z. performed, analysed and interpreted molecular dynamics simulations. D.P: contributed to MEF cell electroporation experiments. S.L: performed cloning and protein purification and assisted with experiments.

